# Internal Transcribed Spacers as Phylogenetic Markers Enable Species-level Metataxonomic Analysis of Ciliated Protozoa

**DOI:** 10.1101/2024.02.02.578691

**Authors:** Sripoorna Somasundaram, Zhongtang Yu

## Abstract

**Background:** The conventional morphology-based classification of ciliates is often inaccurate and time-consuming. To address this issue, sequencing, and analysis of the 18S rRNA gene of ciliates have been used as an alternative. However, this method has limitations because the highly conserved nature of this gene makes it challenging to achieve species-level resolution. This study assesses the capability of two internal transcribed spacers, ITS1 and ITS2, along with the 28S rRNA gene, to enhance the taxonomic resolution beyond that offered by the 18S rRNA gene in free-living and host-associated ciliates.

**Results:** We compared sequences of ITSI, ITS2, and the 18S and the 28S rRNA genes downloaded from public databases and found that ITS1 and ITS2 are more divergent at both inter- and intra-specific levels than the 18S rRNA gene. We designed universal primers specific to the two ITS regions and the 28S rRNA gene for free-living and rumen ciliates. We then systematically evaluated these primers using *in*-*silico* analysis, PCR assays, and metataxonomic or metabarcoding analysis and compared them to universal 18S rRNA gene primers. We found that the new primers are specific and inclusive, with an inclusiveness rate of over 80% based on *in*-*silico* analysis and confirmed their specificity using PCR evaluation. We validated the new primers with metagenomic DNA from freshwater samples and from rumen samples. Our metataxonomic analysis demonstrated that the ITS regions and the 28S rRNA gene could reveal greater ciliate diversity than the 18S rRNA gene in both environments. In particular, ITS1 detected the highest number of ciliate species, including species and genera that were not detected by the 18S rRNA gene.

**Conclusions:** The ITS regions, particularly ITS1, offer superior taxonomic resolution, and the NCBI ITS RefSeq database allows more species to be classified. Therefore, ITS1, and to a lesser extent ITS2, is recommended for enhancing metataxonomic analysis of ciliate communities in both freshwater and rumen environments.

## Background

Microbiomes play pivotal roles in ecosystems, and over the past two decades, significant research efforts have been directed towards microbiome studies in various ecosystems. However, most of the studies have focused primarily on bacteria and archaea, neglecting fungi, ciliates, and viruses. Ciliated protozoa are predators significantly influencing prokaryote populations, and thus profoundly affecting their ecological functions [1]. By preying on or forming symbioses with prokaryotic microbes, ciliates regulate the population dynamics and metabolic processes of prokaryotes, thereby playing an integral role in shaping the structure and the microbial food webs and altering energy transfer across trophic levels in ecosystems [2–4]. Ciliates can either be free-living or associated with other organisms as hosts such as ruminants. Free-living ciliates, having been classified into 12 different classes within the phylum Ciliophora, exhibit remarkable diversity, and can be found in various ecosystems worldwide [5]. In contrast, rumen ciliates only live in the rumen of various ruminants, both wild and domesticated, or similar habitats, and play a crucial role in feed digestion and efficiency, regulation of rumen pH, nitrogen metabolism, and methane emissions [6]. Despite their crucial roles, ciliates remain poorly understood in terms of diversity, population dynamics, and interactions with prokaryotic microbes. This knowledge gap has significant implications for understanding their ecosystem functions [7]. It is thus essential to uncover the often-overlooked diversity of ciliates and investigate their community profiles. The inclusion of ciliates in metataxonomic analysis of microbiomes will support holistic studies of microbiomes.

Traditionally, the identification and classification of ciliates have been solely based on morphological features [8]. However, ciliates have relatively simple morphological features, and these features often provide limited taxonomic resolution at the species level. Moreover, some ciliates cannot be cultured in the laboratory [9], and it requires experience and expertise to distinguish some of the minute diagnostic features [10]. To circumvent these limitations, researchers have used sequencing and analyzing of the small subunit ribosomal RNA (18S rRNA) gene for identification, classification, and community profiling of ciliates [11–13]. However, the highly conserved nature of the 18S rRNA gene limits its capability and utility in species-level taxonomic delimitation among eukaryotes, including ciliates [14, 15]. A few studies have explored alternative phylogenetic markers for analyzing ciliate diversity, such as the 28S rRNA gene [16, 17], *cox*1 [10; 18, 19], *hsp*90 [20], ITS regions [21–24] and *rpo*B [25]. However, the primers designed for these markers are specific to only a few classes or genera of free-living ciliates within the phylum Ciliophora. Furthermore, although genome-resolved metagenomics allows comprehensive analysis of the bacteriomes, archaeomes, and viromes in various microbiomes [26–28], it cannot be efficiently applied to the analysis of ciliate communities due to their unique genome structures (e.g., large genomes, nuclear dimorphism, high ploidy, and chromosomal fragmentation), limited numbers of sequenced ciliate genomes in databases, the lack of a genome taxonomy database, and their relatively low abundance in microbiomes.

Being intrinsically more divergent than the flanking rRNA genes, ITS1 and ITS2 have been widely used in mycobiome (fungal microbiome) analysis for species identification and profiling [29, 30], offering significantly enhanced taxonomic resolution. These two ITS regions have been used in studies of free-living ciliates to infer their evolution and refine their phylogeny among closely related ciliate species [31, 32]. They have also facilitated the discovery of new ciliate species and genera [33–35]. However, their potential for metataxonomic (or metabarcoding) analysis for delineating the diversity and structure of ciliate communities has not been explored. We hypothesize that ITS1, ITS2, and the hypervariable regions of the 28S rRNA gene (hereafter all designated as alternative markers) could provide a finer phylogenetic resolution, thereby enhancing the taxonomic classification of ciliates, both free-living and host-associated. To test this hypothesis, we designed primers specifically targeting each alternative marker (for the 28S rRNA gene, only one hypervariable region with a length suitable for Illumina 2 x 300 paired-end sequencing) of ciliates and evaluated their utility in metataxonomic analysis of freshwater and rumen ciliates. Our findings demonstrate that these alternative phylogenetic markers, particularly ITS1, improve the taxonomic analysis of ciliate communities in freshwater and rumen environments beyond what is achievable using the 18S rRNA gene.

## Methods

### PCR primer design and *in-silico* validation

We conducted a systematic search of public databases to find sequences that span ITS1 and ITS2 of ciliates, along with their flanking the 18S and the 28S rRNA genes and the intervening 5.8S rRNA gene. We found and retrieved approximately 500 DNA sequences of free-living ciliates (both cultured and taxonomically assigned) from GenBank. However, our search yielded only a small number of contiguous sequences in GenBank that encompass the two ITS regions and the rRNA genes of rumen ciliates. We thus created an in-house rRNA operon (18S-ITS1-5.8S-ITS2-28S) database (referred to as rumen ciliate rRNA operon database, RCROD) using the RESCRIPt plugin [36] of QIIME2-2022.8 and the 52 single-cell amplified genomes (SAGs) of rumen ciliates reported by Li et al. (2022) [37]. These SAGs represent 13 genera and 19 distinct species of predominant rumen ciliates (Table S1). We combined the GenBank sequences derived from free-living ciliates and the rumen ciliate sequences, performed alignment using MEGA11 [38], and manually fine-tuned the alignment using BioEdit version 7.2.5 [39]. We then designed primer sets specific to ITS1 and ITS2 using Primer3 [40] based on conserved regions of the 18S, 5.8S, and 28S rRNA genes to enable amplification of the complete ITS regions and maximized primer inclusiveness (Figs. 1 and 2a). Since we could not design universal primers that allow for the amplification of either ITS of both free-living and rumen ciliates, we designed separate primers for free-living ciliates and rumen ciliates (Fig. 1 and Table 1). Similarly, to design primers specifically targeting the 28S rRNA gene, we retrieved approximately 800 28S rRNA gene sequences derived from ciliates from GenBank after excluding those derived from uncultured/unclassified ciliates. After aligning the sequences, we designed one primer set for free-living ciliates and another primer set for rumen ciliates based on the conserved regions that flank the D1-D2 region. The online oligo design tools from ThermoFisher Scientific were used to calculate the GC content and melting temperatures for the designed primers. We did not design primers for the 18S rRNA gene but performed an *in-silico* evaluation of extant primers from the literature for specificity and inclusiveness. We found that the forward primer from Sylvester et al. (2004) [41] and the reverse primer from Ishaq and Wright (2014) [42] have the best inclusiveness and specificity (Table 1). These two primers were used to benchmark the newly designed primers specific to the alternative markers.

To assess the specificity of the newly designed primers, we performed *in*-*silico* PCR using Fast-PCR [43]. Briefly, *in-silico* PCR was conducted with various amplification stringency settings, allowing for fewer than three mismatches per primer-target pairing. The primers that were capable of “amplifying” more than 80% (Fig. 2b) of the respective ciliate sequences were considered inclusive and were synthesized for experimental use.

**Table 1.**
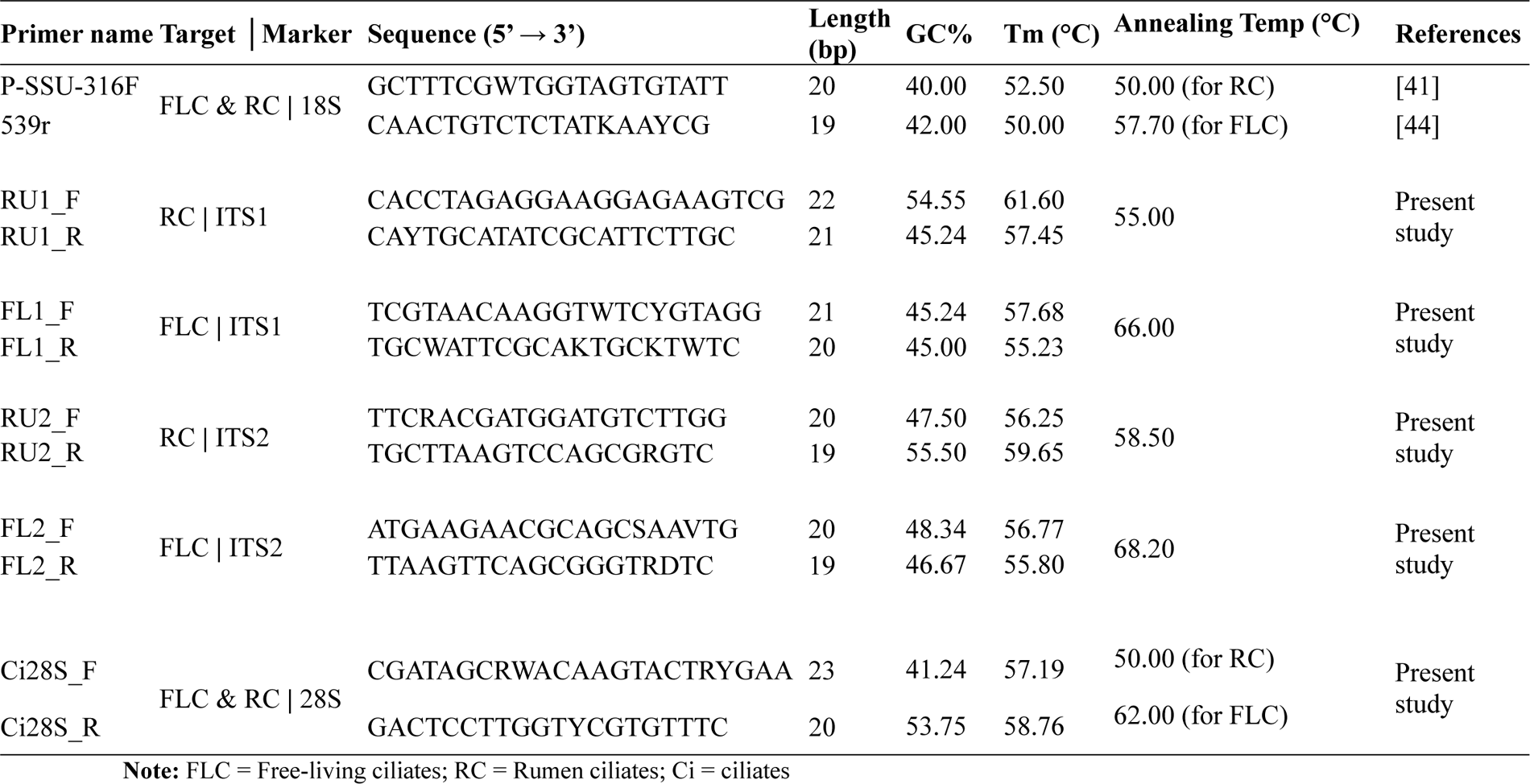
Details of primer sets for amplification of 18S rRNA gene, ITS1, ITS2, and 28S rRNA gene.

**Fig. 1.**
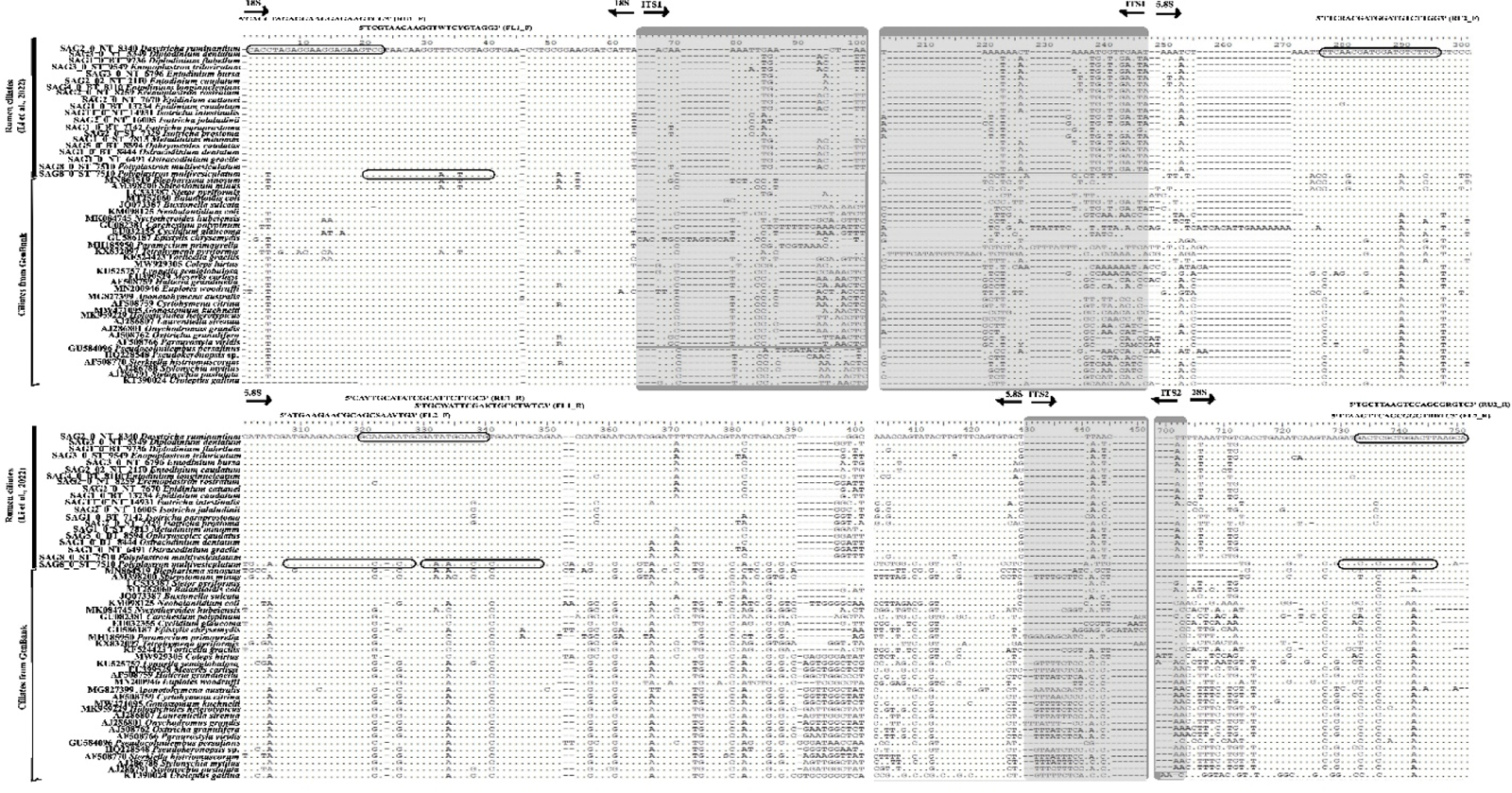
Alignment of ITS sequences of rumen and freshwater ciliates and the locations of the primers. The two sets of primers, i.e., RU1_F and RU1_R, and RU2_F and RU2_R, represent the forward and reverse primers for rumen ciliates for ITS1 and ITS2 regions, respectively. Similarly, FL1_F and RL1_R and FL2_F and RL2_R indicate the forward and reverse primers for freshwater ciliates for ITS1 and ITS2 regions, respectively. The ITS1 and ITS2 regions are shaded in grey.

**Fig. 2.**
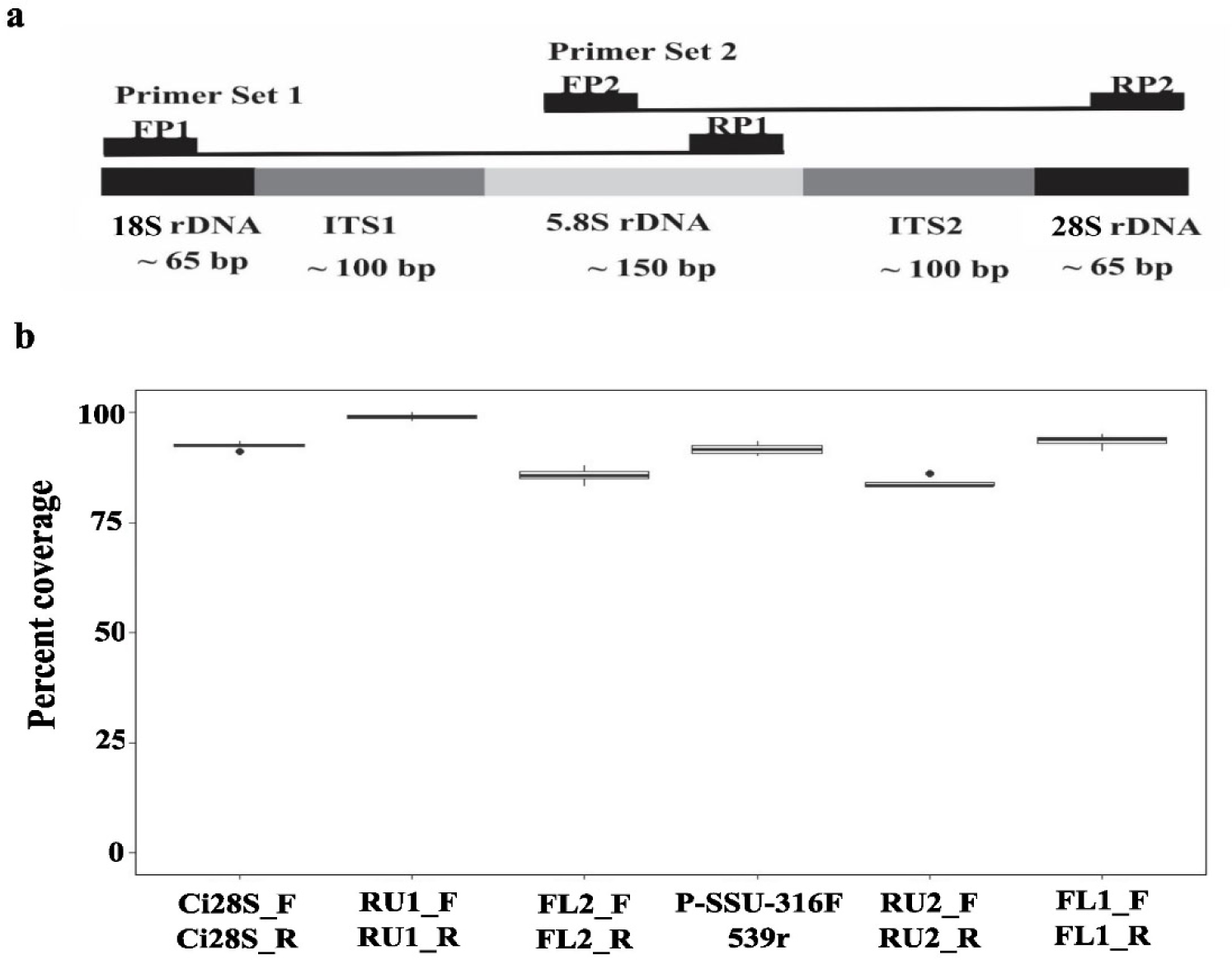
a: A schematic showing the locations of the designed primers to amplify the complete internal transcribed spacer regions, ITS1 and ITS2, and flanking rRNA genes of ciliates; **b:** Comparison of the overall performance (in percent coverage) of the different primer sets used in this study by *in*-*silico* PCR analysis. FP = Forward Primer, RP = Reverse Primer.

### Assessment of the marker sequences for taxonomic resolution

We selected the sequences that span the complete rRNA operons (18S-ITS1-5.8S-ITS2-28S) to assess the taxonomic resolution of each phylogenetic marker. In total, 109 and 66 complete rRNA operons were obtained from free-living and rumen ciliates, respectively. The sequences of each marker (the target region of the 18S and the 28S rRNA genes and the full length of each ITS) were sliced out, combined separately for free-living and rumen ciliates, and aligned using MEGA11. Following manual fine-tuning of the aligned sequences using ClustalX v2 [44], a distance matrix was generated for each marker with the Tamura 3-parameter model applied [45].

The sequence of each marker was also subjected to taxonomic classification using the QIIME2 pipeline [46]. The 18S and the 28S rRNA gene sequences of free-living ciliates were classified using the SILVA_138.1_SSURef_NR99 and SILVA_138.1_LSURef_NR99 databases, respectively. The sequences of ITS1 and ITS2 of free-living ciliates were classified using both the NCBI ITS RefSeq and the UNITE database (https://unite.ut.ee/). The NCBI ITS RefSeq database allowed more ITS1 and ITS2 sequences to be taxonomically assigned compared to the UNITE database, we thus chose the NCBI ITS RefSeq database for further analysis. Since the SILVA, UNITE, and NCBI ITS Ref databases contained very few 18S, ITS1, ITS2, and 28S sequences for rumen ciliates, we classified the sequences of these markers using the RCROD and the genome-based taxonomy of rumen ciliates reported by Li et al. (2022) [37]. The resulting taxonomy tables were converted to Phyloseq objects and then analyzed using the Phyloseq package in R [47]. The total number of ASVs derived from each phylogenetic marker was compared across the four markers.

### Evaluation of each marker for its capability and utility in metataxonomic analysis of freshwater and rumen ciliates

#### Sample collection and processing

Freshwater samples were collected from seven different freshwater bodies in central Ohio, near Columbus, and eight different rumen samples were collected from cannulated dairy cows housed at the Waterman Agricultural and Natural Resources Laboratory of The Ohio State University. The ciliates in the freshwater samples were enriched as described by Gupta et al. (2020) [48] before DNA extraction. Briefly, small pieces of boiled cabbage were added to the freshwater samples in beakers and incubated at 22 °C for 24 h, followed by an additional 48-hour incubation at 22 °C in Petri dishes. The growth of ciliates was microscopically verified. Each sample was then centrifuged at 2,000 rpm for 2 min to pellet the ciliate cells, and the cell pellets were used for DNA extraction. For the rumen samples, each 500 ml sample was strained through four layers of cheesecloth to remove large digesta particles and then mixed with an equal volume of simplex buffer, as outlined by Park et al. (2017) [49]. The mixture was then maintained at 37 °C for 2 hours with continuous CO_2_ sparging to facilitate the removal of small digesta particles. The rumen ciliates were also pelleted by centrifugation at 2,000 rpm for 2 min. The pellets were washed once with the Simplex buffer.

#### DNA extraction and optimization of PCR conditions

Metagenomic DNA was extracted from each pelleted sample using the Qiagen DNeasy Blood and Tissue Kit (Qiagen, Inc., Germantown MD). The quality and quantity of the DNA extracts were evaluated using a Nanodrop spectrophotometer and agarose gel (1%) electrophoresis.

We optimized the PCR conditions for each primer set using gradient PCR, which involved a combination of varying MgCl_2_ concentrations and primer annealing temperatures. The PCR reaction mix (25 µl) contained 1x PCR buffer, 1 µM of each primer, 60 ng of DNA template, 0.32 mM of dNTP (0.08 mM per dNTP), 1.5 mM MgCl_2_, and 0.6 U of Taq DNA polymerase. The PCR thermal cycling conditions were as follows: one initial denaturation step at 95 °C for 5 min; 32 cycles of denaturing at 95 °C for 40 sec, annealing over a range of 50–65 °C (the specific temperature for each primer set is provided in Table 1) for 40 sec, and extension at 72 °C for 40 sec; and final extension step at 72 °C for 10 min.

#### Metataxonomic analyses of freshwater and rumen ciliates

We utilized metataxonomics to evaluate the capability and utility of the new primer sets, in comparison to the 18S rRNA gene primers, in analyzing ciliate communities in both freshwater samples and rumen samples. Briefly, individual amplicon libraries were prepared for each metagenomic DNA sample using each primer set, each synthesized with one distinct unique index sequence for multiplexing. These libraries were prepared following the optimized PCR conditions. After quality checking, the amplicon libraries were pooled at an equal molar ratio and paired-end (2 x 300 bp) sequenced on the Illumina MiSeq system. The sequencing reads were demultiplexed based on the index sequences and further analyzed using the QIIME2 v 2021.4 pipeline. In brief, the primers were trimmed off, and quality plots were generated and visually examined. The regions with low base-calling accuracy were identified using the base quality score (i.e., Phred33). Bases with a quality score <20 were trimmed off using Cutadapt from all the reads before merging the corresponding forward and reverse reads using DADA2 within QIIME2. The merged sequences were then clustered as amplicon sequence variants (ASVs). As previously mentioned, for the freshwater ciliates, the SILVA databases were used to taxonomically classify the ASVs derived from the 18S and the 28S rRNA genes, while the NCBI ITS RefSeq database was used to classify the ITS1 and ITS2 ASVs. The rumen ciliate ASVs of ITS1, ITS2, the 18S, and the 28S rRNA genes were classified with the RCROD database.

#### Alpha diversity estimation and statistical analyses

Prior to estimating alpha diversity, we assessed the level of sample saturation using rarefaction analysis in R. Alpha diversity metrics including Shannon and Simpson diversity indices, species richness (observed and Chao1 estimate), and evenness were calculated using R version 4.2.3 [50]. Statistical analyses were performed using the Mann-Whitney U test (non-parametric) and one-way ANOVA, followed by a post hoc Tukey test (parametric) in R to assess whether the alternative makers could significantly improve the detection of ciliate diversity in comparison to the 18S rRNA gene. We then visualized the data using the ggplot2 [51], tidyverse [52], and vegan [53] packages in R.

## Results and discussion

The diversity of ciliates in various microbiomes remains insufficiently characterized, particularly at the species level, even when analyzed by sequencing their 18S rRNA gene sequences [54]. This is particularly true for rumen ciliates, which are closely related taxa [55, 56]. To overcome this limitation, we evaluated three alternative markers known to have higher sequence divergence among species than the 18S rRNA gene. Utilizing the newly available single-cell amplified genomes of rumen ciliates and the sequences of free-living ciliates in the SILVA and NCBI ITS RefSeq database, we evaluated these markers for sequence variability and compared their sequence dissimilarities to delineate ciliate species. We then designed new inclusive primers targeting these markers and confirmed their specificity. These alternative markers, when amplified with the new primers, produced amplicons of an appropriate length for metataxonomic analysis of free-living and rumen ciliates, and improved species-level resolution compared to the 18S rRNA gene.

### The database sequences of the alternative markers are more divergent than the 18S rRNA gene, potentially providing a higher taxonomic resolution

The *in-silico* analysis of database sequences revealed that the alternative markers share a lower similarity than the 18S rRNA gene, both among the selected 109 free-living ciliate sequences and among the 66 rumen ciliate sequences (Table 2). This is expected and aligns with previous findings among prokaryotic and other eukaryotic microbes [57, 58]. Across the free-living ciliates, the sequence similarities of the ITS regions and 28S rRNA gene varied from 10 to 19%, whereas that of the 18S rRNA gene reached approximately 64%. Similarly, the sequence similarity for ITS1 and the 28S rRNA gene of rumen ciliates was also much lower than that of the 18S rRNA gene (Table 2). Among both free-living and rumen ciliates, ITS1 exhibited very low sequence similarity, approximately 18% among free-living ciliates and 12% among rumen ciliates. However, all the markers, except for ITS1, showed a higher sequence similarity among rumen ciliates when compared to free-living ciliates (Table 2), corroborating their limited taxonomic distribution within the class Litostomatea. Compared to the rRNA genes, the ITS regions experience a lower level of mutational constraint, allowing for increased sequence variability [59]. This increased sequence variability of the alternative markers potentially enhances the taxonomic resolution of sequence-based analysis of ciliate communities.

Sequence analysis of the four markers of 109 free-living ciliates and the 66 rumen ciliates revealed that each marker sequence represented one ASV. More ASVs derived from ITS1 and ITS2 sequences were assigned to known species than those derived from the 18S and the 28S rRNA gene sequences (Table 2). Importantly, in contrast to the two rRNA genes, a greater number of species were identified from the ASVs derived from the ITS1 sequences of both free-living and rumen ciliates (Table 2). Additionally, more known orders, families, and genera of both free-living and rumen ciliates were identified by the ASVs derived from ITS1 and ITS2. Compared to the ASVs derived from the 18S rRNA gene sequences, fewer ASVs derived from the 28S rRNA gene sequences were assigned to genera or species, which is likely due to the smaller number of 28S rRNA gene sequences than the 18S rRNA gene sequences in the SILVA Ref database (24,708 vs.167,330). Collectively, the sequences of the two ITS markers enhanced the taxonomic classification of the ASVs at the order, family, genus, and species levels, beyond that achievable based on the 18S and the 28S rRNA gene sequences. The ASVs derived from sequences of ITS1 and ITS2 also captured a wide array of known taxa including genera and species. These findings demonstrate that both ITS regions, particularly ITS1, can enhance the detection and identification of both free-living and rumen ciliates.

**Table 2.** Sequence* similarities; numbers of ASVs assigned to known genera and species; numbers of known taxa to which the ASVs could be assigned; and numbers of known species exclusively “detected” by each marker.

| Ciliates | Markers | Sequence similarity (%) | Genus-level ASVs | Species-level ASVs | Order | Family | Genus | Species | Marker-exclusive species |
| --- | --- | --- | --- | --- | --- | --- | --- | --- | --- |
| Free-living | 18S | 64.04 | 57 | 9 | 1 | 0* | 6 | 2 | 1 |
|  | ITS1 | 18.18 | 88 \| § 43 | 85 \| 47 | 6 \| 4 | 15 \| 6 | 15 \| 7 | 20 \| 10 | 8 \| 1 |
|  | ITS2 | 09.94 | 93 \| 56 | 78 \| 57 | 6 \| 5 | 10 \| 9 | 14 \| 10 | 16 \| 12 | 5 \| 1 |
|  | 28S | 18.67 | 38 | 29 | 0 | 0 | 3 | 4 | 2 |
| Rumen | 18S | 79.24 | 50 | 50 | 2 | 2 | 10 | 11 | 0 |
|  | ITS1 | 11.57 | 64 | 60 | 2 | 2 | 10 | 16 | 3 |
|  | ITS2 | 54.81 | 53 | 40 | 2 | 2 | 7 | 8 | 0 |
|  | 28S | 31.79 | 3 | 3 | 1 | 1 | 1 | 1 | 0 |

\* The sequences for free-living ciliates were downloaded from NCBI ITS RefSeq (109 sequences, each representing one ASV) that encompass the complete operons (18S-ITS1-5.8S-ITS2-28S rRNA), and the sequences for rumen protozoa (66 sequences, each also representing one ASV) were from one custom database containing 52 single-cell-amplified genomes reported by Li et al. (2022).
\* The lack of information in the SILVA/NCBI ITS RefSeq databases used for the current investigation could be the reason for the '0' values of orders and families detected by the 18S and 28S rRNA gene sequences.
§ For free-living ciliates, the values before and after the “ | ” were based on the NCBI ITS RefSeq database and the UNITE database, respectively. The ITS sequences of rumen ciliates were only analyzed with the custom database.

Some of the genera represented by the database sequences encompassed multiple species, including closely related ones (phylogenetic distance = 0 or 0.01 based on the 18S gene sequences). To determine if the alternative markers can differentiate the closely related species, we compared the pair-wise distance matrices derived from these markers. For free-living ciliates, both ITS regions were able to differentiate between these closely related species pairs, with the distance values ranging from 0.15 to 0.61 for ITS1 and 0.04 to 0.23 for ITS2, except for one species pair (*Holostrichides heterotypicus* with *Holostrichides obliquocirratus*), which was not adequately differentiated by ITS2 (with a distance matrix value of 0.01). In contrast, the 28S rRNA gene was unable to differentiate two of the species pairs (phylogenetic distance = 0), while the rest were differentiated with small distance matrix values (Table 3). For the rumen ciliates, the ITS regions and 28S rRNA gene were able to distinguish between all the closely related species pairs (distance matrix values ranging from 0.02 to 0.05) except for two species pairs (*Entodinium bursa* and *Entodinium caudatum,* and *Isotricha jalaludinii* and *Isotricha intestinalis*) not being distinguished by ITS2 and one species pair (*En. bursa* with *En. caudatum*) not being distinguished by the 28S rRNA gene. In summary, the highest pair-wise distance between closely related species was observed from ITS1, followed by ITS2 and the 28S rRNA gene. These findings are consistent with those reported for five astome ciliate species [24] and demonstrate that ITS1 can differentiate closely related ciliate species better than the 18S rRNA gene.

**Table 3.** Phylogenetic distance (mean±SD) between two select species that each have multiple database sequences.

| Species | 18S | ITS1 | ITS2 | 28S |
| --- | --- | --- | --- | --- |
| **Free-living ciliates** |  |  |  |  |
| *Holotrichides heterotypicus* vs *H. obliquocirratus* | 0.00±0.00* | 0.60±0.00<sup>a</sup> | 0.01±0.00** | 0.00±0.00* |
| *Spirostomum dharwarensis* vs *S. ambiguum* | 0.01±0.00** | 0.61±0.37<sup>b</sup> | 0.15±0.10 | 0.01±0.00** |
| *Spirostomum dharwarensis* vs *S. minus* | 0.01±0.00** | 0.57±0.36<sup>c</sup> | 0.08±0.05 | 0.02±0.01 |
| *Spirostomum dharwarensis* vs *S. subtilis* | 0.01±0.00** | 0.51±0.49 | 0.15±0.10 | 0.01±0.00** |
| *Spirostomum minus* vs *S. ambiguum* | 0.01±0.00** | 0.49±0.45<sup>a</sup> | 0.23±0.05<sup>a</sup> | 0.01±0.00** |
| *Spirostomum minus* vs *S. subtilis* | 0.01±0.00** | 0.15±0.01<sup>a</sup> | 0.05±0.01 | 0.01±0.01** |
| *Spirostomum teres* vs *S. teres* | 0.00±0.00* | 0.44±0.31<sup>a</sup> | 0.04±0.02 | 0.02±0.01 |
| *Stentor pyrififormis* vs *Stentor amethystinus* | 0.00±0.00* | 0.51±0.21<sup>a</sup> | 0.12±0.06<sup>c</sup> | 0.00±0.00* |
| **Rumen ciliates** |  |  |  |  |
| *Entodinium bursa* vs *E. caudatum* | 0.00±0.00* | 0.03±0.02<sup>b</sup> | 0.01±0.00** | 0.01±0.00** |
| *Entodinium bursa* vs *E. loginucleatum* | 0.00±0.00* | 0.02±0.01 | 0.03±0.02 | 0.04±0.02<sup>c</sup> |
| *Isotricha jalaludinii* vs *I. intestinalis* | 0.00±0.00* | 0.02±0.01 | 0.00±0.00* | 0.05±0.03<sup>b</sup> |
| *Isotricha jalaludinii* vs *I. paraprostoma* | 0.00±0.00* | 0.04±0.02<sup>c</sup> | 0.04±0.01 | 0.04±0.02<sup>c</sup> |
| *Isotricha prostoma* vs *I. jalaludinii* | 0.00±0.00* | 0.04±0.03 | 0.04±0.02 | 0.03±0.02 |

**Note:** Distance was calculated with the Tamura 3-parameter model applied. 0 and 0.01 distances are represented by \* and \*\*, respectively.
The superscripts indicate the level of significance when compared to the 18S rRNA gene: a, $p < 0.01$ ; b, $p = 0.01$ ; c, $p < 0.05$ .

### The alternative markers improve metataxonomic analysis of freshwater and rumen ciliates

#### Summary of sequencing data

The new primers designed for ITS1, ITS2, and the 28S rRNA gene all successfully amplified their respective targets, yielding amplicons ranging from 230 to 440 bp. Since the resultant amplicons are all shorter than 450 bp, they are of appropriate lengths for sequencing using the Illumina MiSeq system, which is the primary sequencing technology used for metataxonomic analysis.

We subsequently compared the four markers for their capability to detect and identify ciliates using freshwater and rumen samples. Following filtering, denoising, and removal of sequence chimera, we obtained a total of 415,688 sequences of the 18S rRNA gene, 792,360 sequences of ITS1, 308,212 sequences of ITS2, and 241,117 sequences of the 28S rRNA gene from the rumen samples. The freshwater samples yielded a total of 17,706 sequences of the 18S rRNA gene, 157,706 ITS1 sequences, 306,636 ITS2 sequences for ITS2, and 93,578 sequences of the 28S rRNA gene. The number of quality-checked sequences ranged from 2,000 to 89,000 per sample for each marker.

#### Richness and diversity of ciliates detected in freshwater and rumen samples

The alternative markers generated more ASVs from both the freshwater and the rumen samples, both in total and per sample (Fig. 3a-3f). Statistical analyses revealed that all three alternative markers yielded a significantly greater number of ASVs than the 18S rRNA gene from the freshwater samples (p < 0.05). From the rumen samples, ITS1 and the 28S rRNA gene, but not ITS2, yielded a greater number of ASVs (p < 0.05) compared to the 18S rRNA gene.

**Fig 3.**
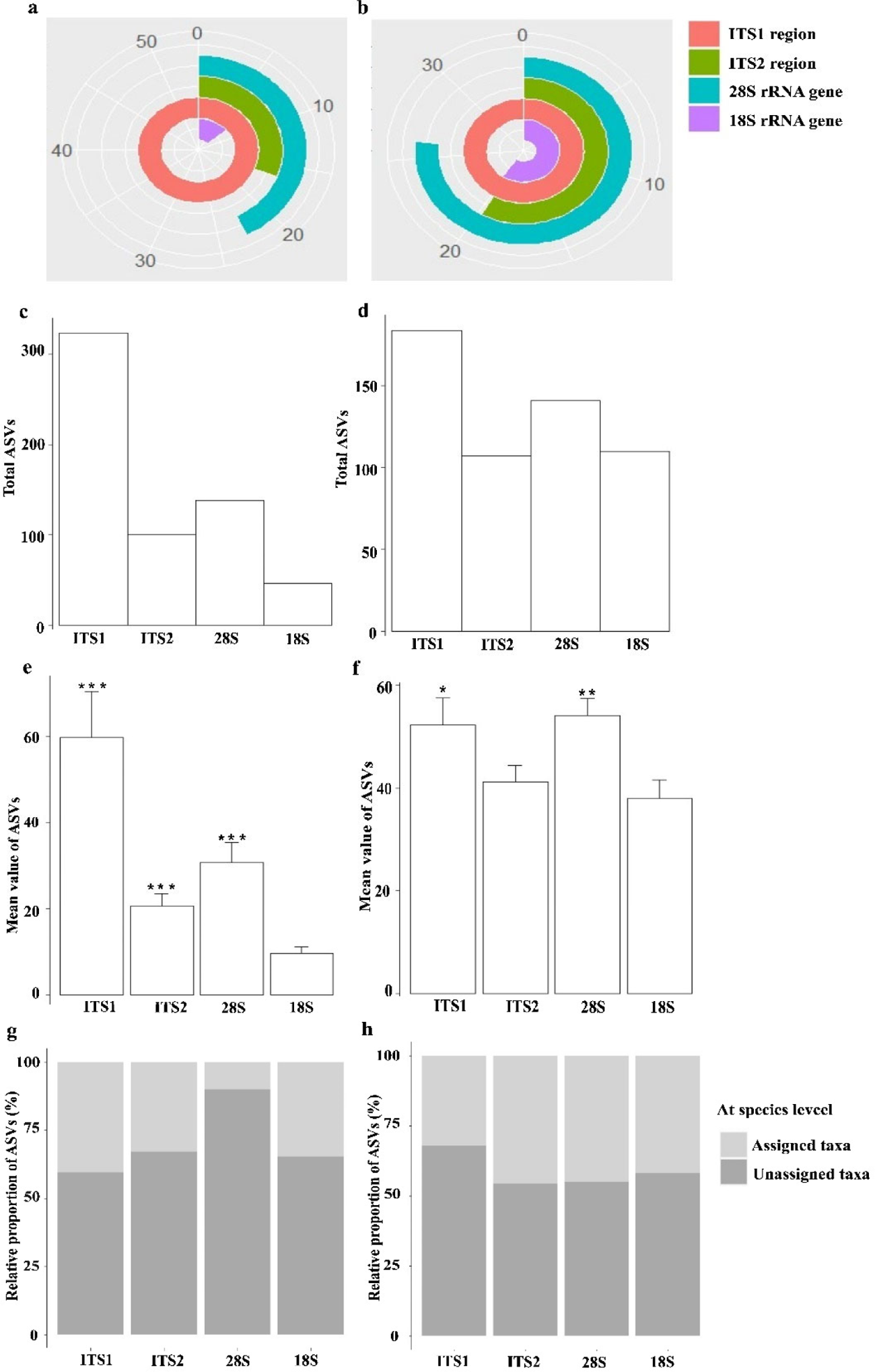
Taxonomic richness determined by the four phylogenetic markers in the freshwater (**a, c, e, and g**) and the rumen (**b, d, f, and h**) samples. **a, b:** total number of ASVs; **c, d:** number of ASVs per sample; **e, f:** mean numbers of ASVs; **g, h:** percentage of assigned and unassigned taxa at the species level. Statistical significance was tested using one-way ANOVA and Mann-Whitney U test: ***, p<0.01; **, p=0.01; *, p < 0.05.

Analysis of alpha diversity revealed significantly greater (*P* <0.05) diversity within freshwater ciliate communities when assessed using ITS1 and the 28S rRNA gene than using the 18S rRNA gene, as indicated by both Shannon diversity index and species richness (both observed and the Chao1 estimate) (Fig. 4). This trend was also observed across the rumen ciliate communities, except for ITS1 with respect to diversity indices (Shannon and Simpson index), which is only numerically greater compared to the 18S rRNA gene. However, in the freshwater ciliate communities, ITS2 revealed greater richness (both observed and estimated) than the 18S rRNA gene, while maintaining comparable diversity indices. The greater diversity indices and species richness of rumen ciliates detected using ITSI than ITS2 align with the much lower sequence similarity of the former than the latter (Table 2). The heightened alpha diversity captured by ITS1 and the 28S rRNA gene aligns with their elevated sequence variability. This suggests that ITS1, and to a lesser extent the 28S rRNA, can improve diversity analysis of ciliate communities in both freshwater and the rumen samples.

**Fig. 4.**
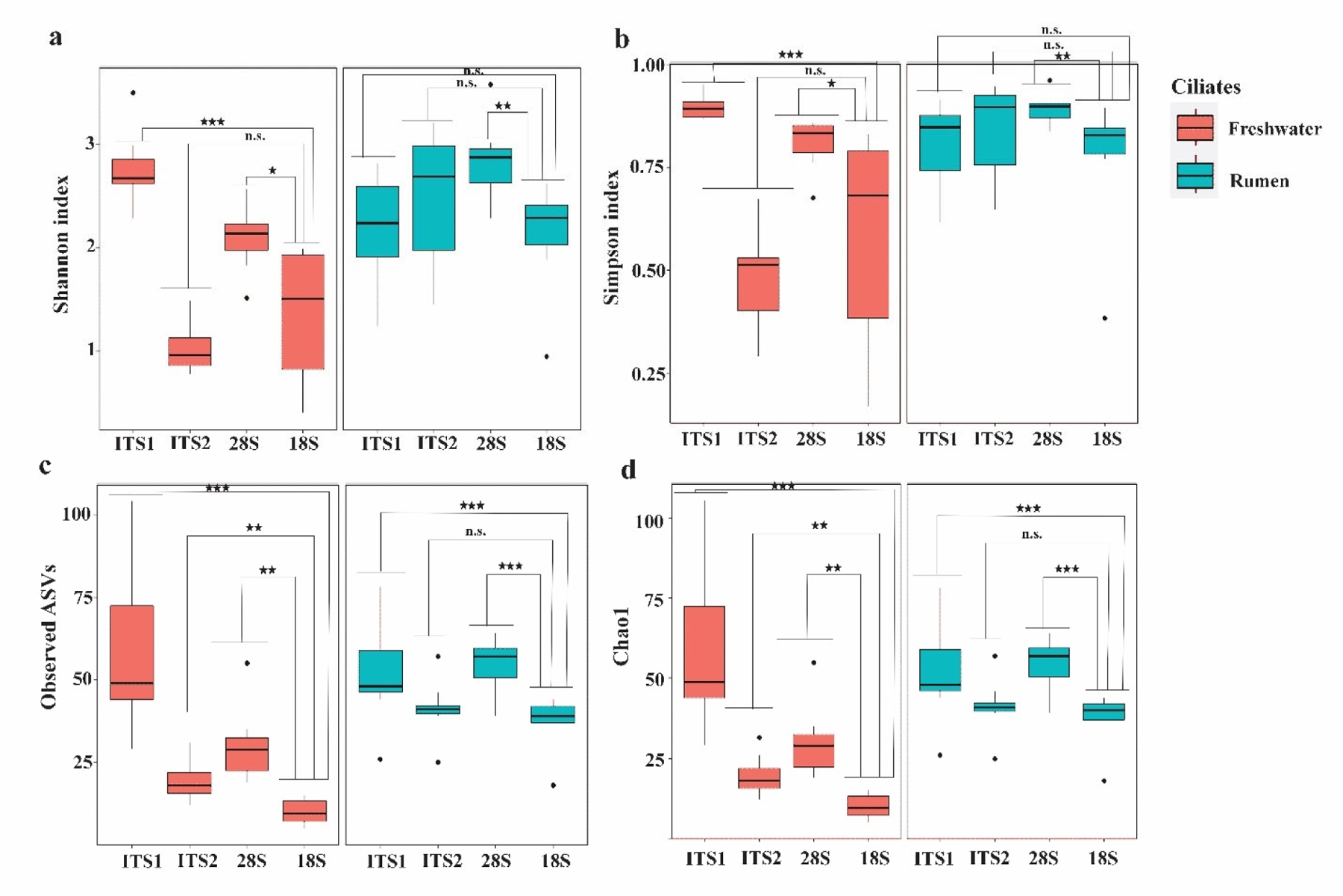
Alpha diversity indices of freshwater and rumen ciliates. Statistical significance was tested using one-way ANOVA and Mann-Whitney U test. ***, p<0.01; **, p=0.01; *, p < 0.05; and n.s., not significant.

#### Ciliates species and higher taxa detected and identified

We compared the percentages of ASVs that were classified to known taxa (from class to species) among the phylogenetic markers. Within the freshwater samples, approximately only 10% of the ASVs detected by the 28S rRNA gene were classified, while a higher percentage of the ASVs detected by the other three markers were classified (Fig. 3g). In the rumen samples, the percentages of assigned and unassigned ASVs were similar across all the four phylogenetic markers, except for ITS1 showing a relatively lower percentage of assigned ASVs. However, despite the lower percentage of classified ASVs derived from ITS1 in the rumen samples and the 28S rRNA gene in the freshwater samples, these two markers detected and identified a greater number of total ASVs.

Within the freshwater samples, ITS1, followed by ITS2, detected a greater number of classes, orders, families, genera, and species than the two rRNA genes (Table 4). The 18S and the 28S genes did not detect any orders or families, except for one order detected by the 18S rRNA gene. Such a poor detection of classified taxa of freshwater ciliates at these taxonomic ranks can be attributed to the limited numbers of classified ciliates recorded in the SILVA database though PCR bias might also be a contributing factor. To validate this premise, we searched the SILVA database for the classes, orders, and families of free-living ciliates recorded in the NCBI ITS RefSeq database. Although the SILVA_138.1_SSURef_NR99 and SILVA_138.1_LSURef_NR99 databases contain some subclasses and suborders of protozoa, they individually represent a much smaller number of classes, orders, and families of ciliate than the NCBI ITS RefSeq database. Notably, 11 classes, nine orders, and two families of ciliates were recorded in SILVA_138.1_SSURef_NR99, whereas SILVA_138.1_LSURef_NR99 has only four classes without any orders and one family recorded. The NCBI ITS RefSeq database contained many more classified taxa: 11 classes, 34 orders, and 108 families. A further comparison revealed that SILVA_138.1_SSURef_NR99 and NCBI ITS RefSeq had the same set of 11 ciliate classes (Litostomatea, Armophorea, Plagiopylea, Oligohymenophorea, Phyllopharyngea, Spirotrichea, Nassphorea, Karyorelictea, Colpodea, Prostomatea, and Heterotrichea), while SILVA_138.1_LSURef_NR99 contained only four classes (Litostomatea, Prostomatea, Oligohymenophorea, and Spirotrichea) that were found in the other two databases. Furthermore, only four orders (Clevelandellida, Colpodida, Cyrtolophosididia, and Plagiopylida) were found to be common between SILVA_138.1_SSURef_NR99 and NCBI ITS RefSeq. At the family level, only Oxytrichidae and Mesodiniidae were found in both SILVA_138.1_SSURef_NR99 and NCBI ITS RefSeq, and Mesodiniidae is the only family documented across all three databases. These results, therefore, indicate that NCBI ITS RefSeq has more classified taxa, particularly at the class, order, and family levels, and a better-defined taxonomy of ciliates than the two SILVA databases, which can improve metataxonomic analysis of ciliate communities.

In the freshwater samples, the 18S rRNA gene detected 13 ciliate species, while ITS1 detected 29 ciliate species (Tables 4 and 5). Of the species detected by ITS1, 26 were not detected by the 18S rRNA gene. ITS2 also detected more species, 20 in total, than the 18S rRNA gene, with 17 of them not being detected by the 18S rRNA gene. However, both ITS1 and ITS2 did not detect 10 of the species detected by the 18S rRNA gene (Table 5). The 28S rRNA gene detected the fewest ciliate species (4 in total) in the freshwater samples. As explained above, the limited records of ciliate species in the current SILVA database might be one reason for the smaller number of ciliate species detected by both rRNA gene markers. Thirteen freshwater ciliate species were detected by both ITS1 and ITS2, leaving 16 species detected exclusively by ITS1 and seven species only detected by ITS2 (Table 5). Thus, while ITS1 and ITS2 could detect a large number of freshwater ciliate species, a small number of ciliate species may escape detection by these two ITS markers. Out of the 47 freshwater ciliate species detected by two or three of the four markers, most of them belong to the classes Oligohymenophorea and Spirotrichea (15 and 25 species, respectively, Table 5). Further, some ciliate species belonging to the classes Heterotrichea, Phyllopharyngea, and Prostomatea were identified only by the ITS region (Table 5, Fig. 5a). A combination of two or more markers, particularly ITS1 and the 18S rRNA gene, may be used together to enhance comprehensive metataxonomic analysis of ciliate communities in freshwater ecosystems.

**Table 4.**
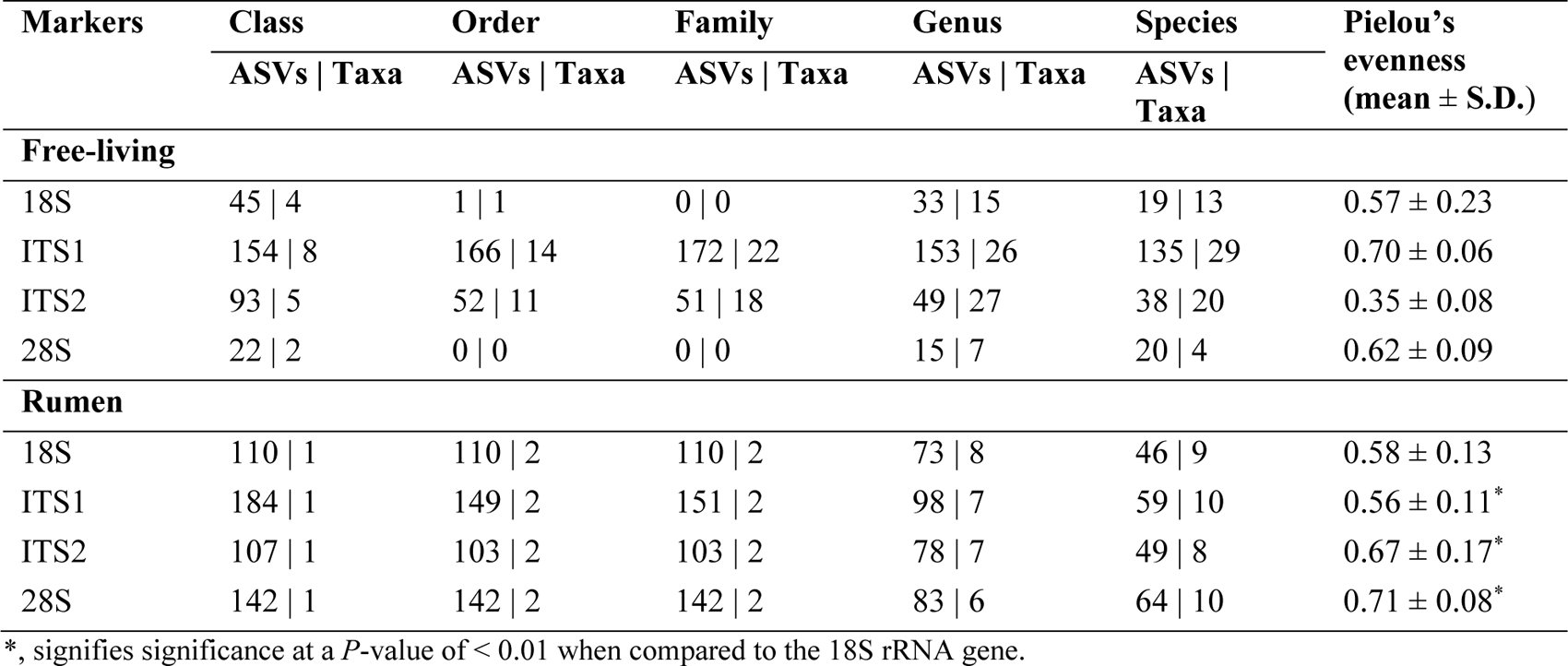
Numbers of ASVs | number taxa assigned at different taxonomic levels; and species evenness detected by each phylogenetic marker.

| Markers | Class | Order | Family | Genus | Species | Pielou's evenness (mean ± S.D.) |
| --- | --- | --- | --- | --- | --- | --- |
|  | ASVs \| Taxa | ASVs \| Taxa | ASVs \| Taxa | ASVs \| Taxa | ASVs \| Taxa |  |
| Free-living |  |  |  |  |  |  |
| 18S | 45 \| 4 | 1 \| 1 | 0 \| 0 | 33 \| 15 | 19 \| 13 | 0.57 ± 0.23 |
| ITS1 | 154 \| 8 | 166 \| 14 | 172 \| 22 | 153 \| 26 | 135 \| 29 | 0.70 ± 0.06 |
| ITS2 | 93 \| 5 | 52 \| 11 | 51 \| 18 | 49 \| 27 | 38 \| 20 | 0.35 ± 0.08 |
| 28S | 22 \| 2 | 0 \| 0 | 0 \| 0 | 15 \| 7 | 20 \| 4 | 0.62 ± 0.09 |
| Rumen |  |  |  |  |  |  |
| 18S | 110 \| 1 | 110 \| 2 | 110 \| 2 | 73 \| 8 | 46 \| 9 | 0.58 ± 0.13 |
| ITS1 | 184 \| 1 | 149 \| 2 | 151 \| 2 | 98 \| 7 | 59 \| 10 | 0.56 ± 0.11* |
| ITS2 | 107 \| 1 | 103 \| 2 | 103 \| 2 | 78 \| 7 | 49 \| 8 | 0.67 ± 0.17* |
| 28S | 142 \| 1 | 142 \| 2 | 142 \| 2 | 83 \| 6 | 64 \| 10 | 0.71 ± 0.08* |

\*, signifies significance at a $P$ -value of $< 0.01$ when compared to the 18S rRNA gene.

**Table 5.**
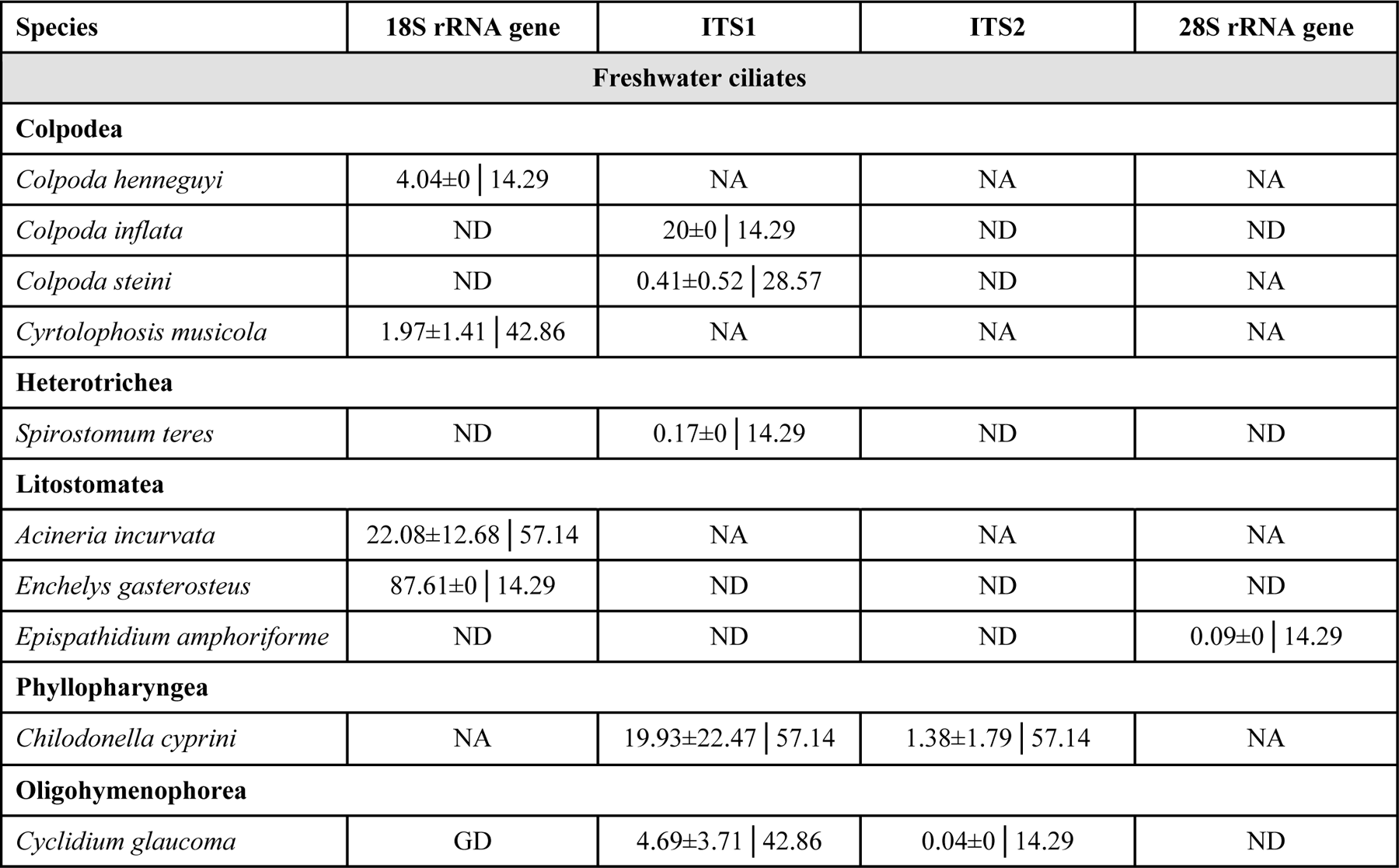
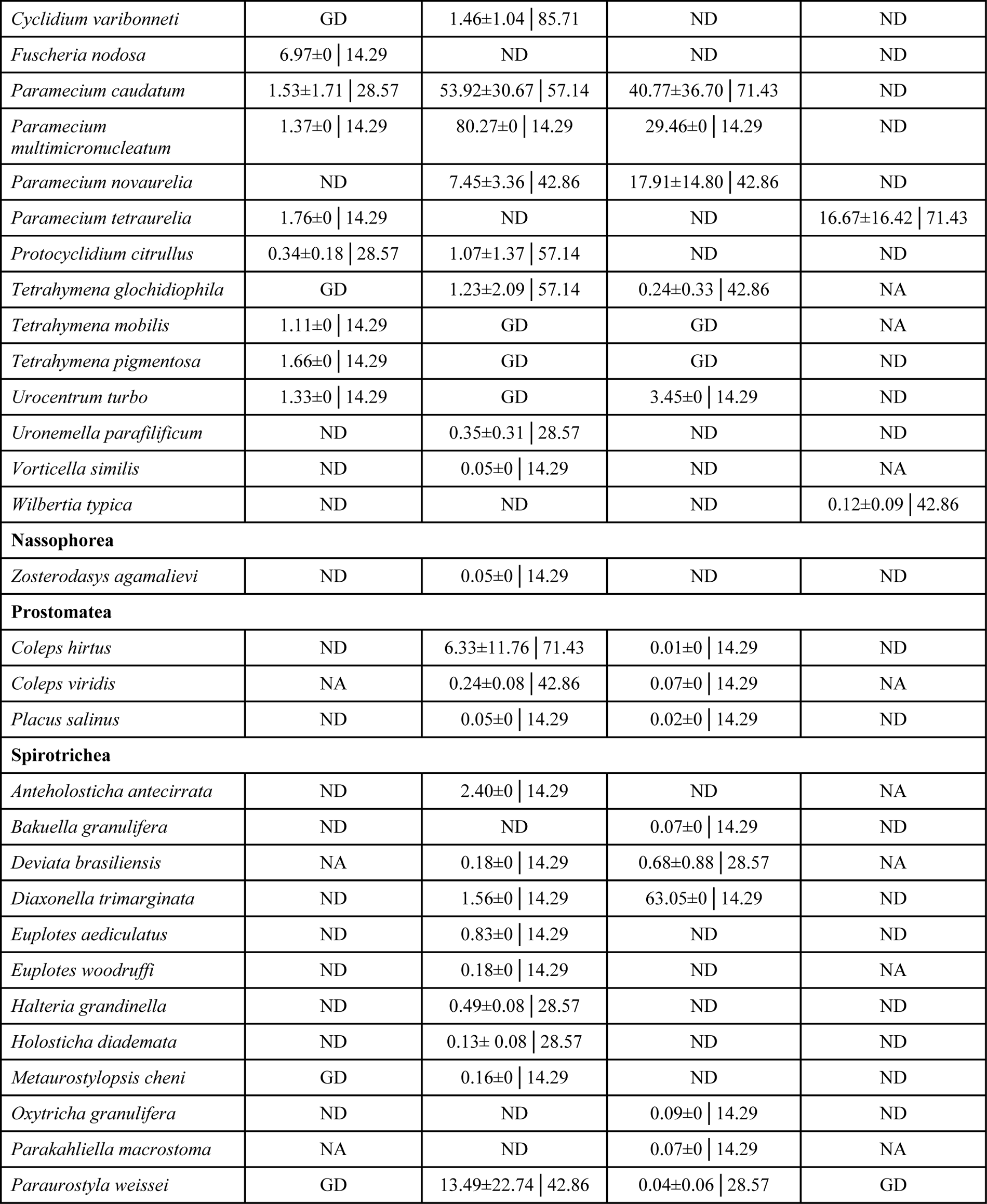
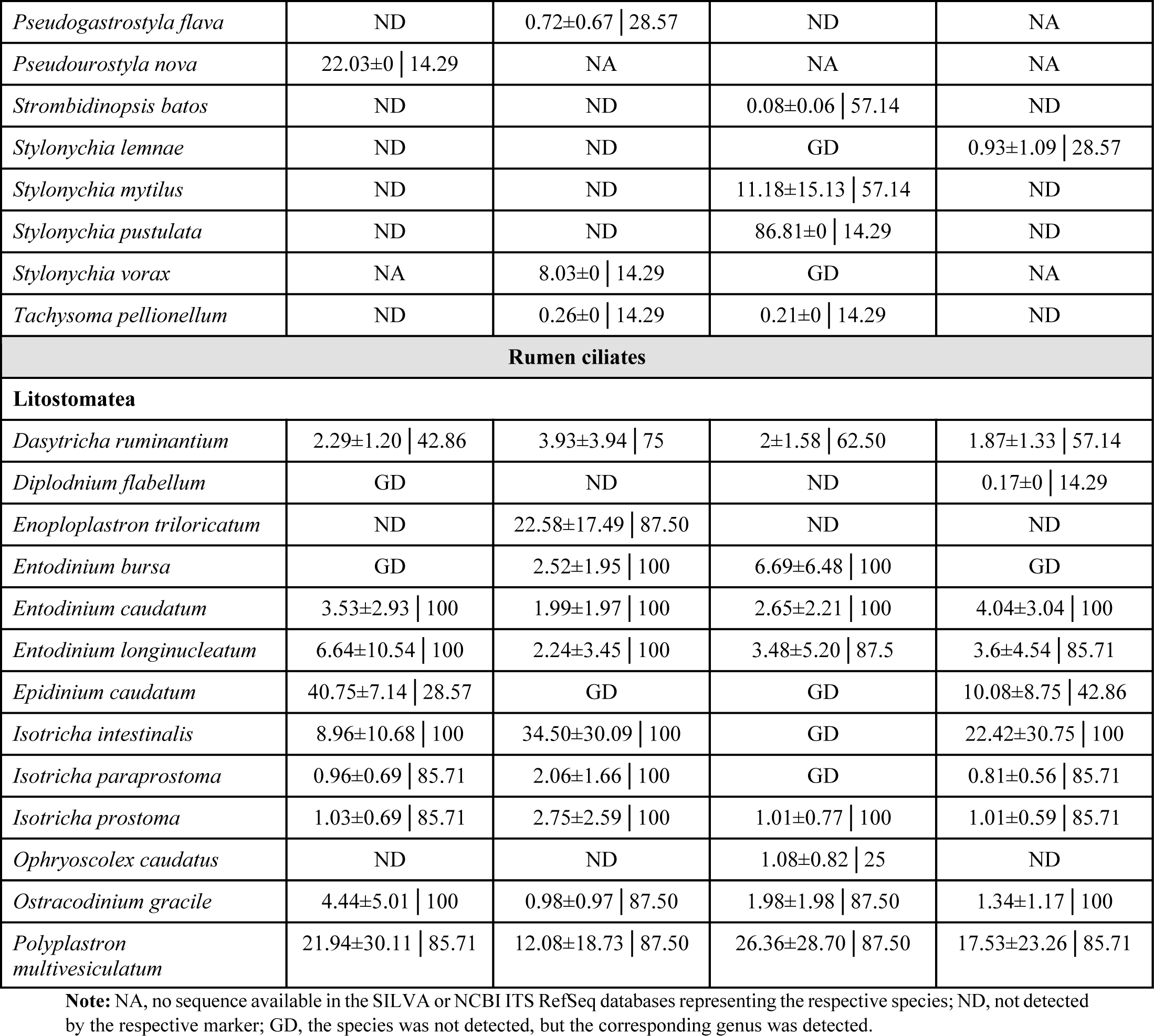
Relative abundance (% ± SD)│prevalence (%) of the identified ciliate species (listed per class) detected by the four phylogenetic markers.

| Species | 18S rRNA gene | ITS1 | ITS2 | 28S rRNA gene |
| --- | --- | --- | --- | --- |
| **Freshwater ciliates** |  |  |  |  |
| **Colpodea** |  |  |  |  |
| *Colpoda henneguyi* | 4.04±0 \| 14.29 | NA | NA | NA |
| *Colpoda inflata* | ND | 20±0 \| 14.29 | ND | ND |
| *Colpoda steini* | ND | 0.41±0.52 \| 28.57 | ND | NA |
| *Cyrtolophosis musicola* | 1.97±1.41 \| 42.86 | NA | NA | NA |
| **Heterotrichea** |  |  |  |  |
| *Spirostomum teres* | ND | 0.17±0 \| 14.29 | ND | ND |
| **Litostomatea** |  |  |  |  |
| *Acineria incurvata* | 22.08±12.68 \| 57.14 | NA | NA | NA |
| *Enchelys gasterosteus* | 87.61±0 \| 14.29 | ND | ND | ND |
| *Epispathidium amphoriforme* | ND | ND | ND | 0.09±0 \| 14.29 |
| **Phyllopharyngea** |  |  |  |  |
| *Chilodonella cyprini* | NA | 19.93±22.47 \| 57.14 | 1.38±1.79 \| 57.14 | NA |
| **Oligohymenophorea** |  |  |  |  |
| *Cyclidium glaucoma* | GD | 4.69±3.71 \| 42.86 | 0.04±0 \| 14.29 | ND |
| *Cyclidium varibonneti* | GD | 1.46±1.04 \| 85.71 | ND | ND |
| *Fuscheria nodosa* | 6.97±0 \| 14.29 | ND | ND | ND |
| *Paramecium caudatum* | 1.53±1.71 \| 28.57 | 53.92±30.67 \| 57.14 | 40.77±36.70 \| 71.43 | ND |
| *Paramecium multimicronucleatum* | 1.37±0 \| 14.29 | 80.27±0 \| 14.29 | 29.46±0 \| 14.29 | ND |
| *Paramecium novaurelia* | ND | 7.45±3.36 \| 42.86 | 17.91±14.80 \| 42.86 | ND |
| *Paramecium tetraurelia* | 1.76±0 \| 14.29 | ND | ND | 16.67±16.42 \| 71.43 |
| *Protocyclidium citrullus* | 0.34±0.18 \| 28.57 | 1.07±1.37 \| 57.14 | ND | ND |
| *Tetrahymena glochidiophila* | GD | 1.23±2.09 \| 57.14 | 0.24±0.33 \| 42.86 | NA |
| *Tetrahymena mobilis* | 1.11±0 \| 14.29 | GD | GD | NA |
| *Tetrahymena pigmentosa* | 1.66±0 \| 14.29 | GD | GD | ND |
| *Urocentrum turbo* | 1.33±0 \| 14.29 | GD | 3.45±0 \| 14.29 | ND |
| *Uronemella parafilificum* | ND | 0.35±0.31 \| 28.57 | ND | ND |
| *Vorticella similis* | ND | 0.05±0 \| 14.29 | ND | NA |
| *Wilbertia typica* | ND | ND | ND | 0.12±0.09 \| 42.86 |
| **Nassophorea** |  |  |  |  |
| *Zosterodasys agamalievi* | ND | 0.05±0 \| 14.29 | ND | ND |
| **Prostomea** |  |  |  |  |
| *Coleps hirtus* | ND | 6.33±11.76 \| 71.43 | 0.01±0 \| 14.29 | ND |
| *Coleps viridis* | NA | 0.24±0.08 \| 42.86 | 0.07±0 \| 14.29 | NA |
| *Placus salinus* | ND | 0.05±0 \| 14.29 | 0.02±0 \| 14.29 | ND |
| **Spirotrichea** |  |  |  |  |
| *Anteolosticha antecirrata* | ND | 2.40±0 \| 14.29 | ND | NA |
| *Bakuella granulifera* | ND | ND | 0.07±0 \| 14.29 | ND |
| *Deviata brasiliensis* | NA | 0.18±0 \| 14.29 | 0.68±0.88 \| 28.57 | NA |
| *Diaxonella trimarginata* | ND | 1.56±0 \| 14.29 | 63.05±0 \| 14.29 | ND |
| *Euplotes aediculatus* | ND | 0.83±0 \| 14.29 | ND | ND |
| *Euplotes woodruffi* | ND | 0.18±0 \| 14.29 | ND | NA |
| *Halteria grandinella* | ND | 0.49±0.08 \| 28.57 | ND | ND |
| *Holosticha diademata* | ND | 0.13±0.08 \| 28.57 | ND | ND |
| *Metaurostylopsis cheni* | GD | 0.16±0 \| 14.29 | ND | ND |
| *Oxytricha granulifera* | ND | ND | 0.09±0 \| 14.29 | ND |
| *Parakahliella macrostoma* | NA | ND | 0.07±0 \| 14.29 | NA |
| *Paraurostyla weissei* | GD | 13.49±22.74 \| 42.86 | 0.04±0.06 \| 28.57 | GD |
| *Pseudogastrostyla flava* | ND | 0.72±0.67 \| 28.57 | ND | NA |
| *Pseudourostyla nova* | 22.03±0 \| 14.29 | NA | NA | NA |
| *Strombidinopsis batos* | ND | ND | 0.08±0.06 \| 57.14 | ND |
| *Stylonychia lemnae* | ND | ND | GD | 0.93±1.09 \| 28.57 |
| *Stylonychia mytilus* | ND | ND | 11.18±15.13 \| 57.14 | ND |
| *Stylonychia pustulata* | ND | ND | 86.81±0 \| 14.29 | ND |
| *Stylonychia vorax* | NA | 8.03±0 \| 14.29 | GD | NA |
| *Tachysoma pellionellum* | ND | 0.26±0 \| 14.29 | 0.21±0 \| 14.29 | ND |
| **Rumen ciliates** |  |  |  |  |
| **Litostomatea** |  |  |  |  |
| *Dasytricha ruminantium* | 2.29±1.20 \| 42.86 | 3.93±3.94 \| 75 | 2±1.58 \| 62.50 | 1.87±1.33 \| 57.14 |
| *Diplodnium flabellum* | GD | ND | ND | 0.17±0 \| 14.29 |
| *Enoploplastron triloricatum* | ND | 22.58±17.49 \| 87.50 | ND | ND |
| *Entodinium bursa* | GD | 2.52±1.95 \| 100 | 6.69±6.48 \| 100 | GD |
| *Entodinium caudatum* | 3.53±2.93 \| 100 | 1.99±1.97 \| 100 | 2.65±2.21 \| 100 | 4.04±3.04 \| 100 |
| *Entodinium longinucleatum* | 6.64±10.54 \| 100 | 2.24±3.45 \| 100 | 3.48±5.20 \| 87.5 | 3.6±4.54 \| 85.71 |
| *Epidinium caudatum* | 40.75±7.14 \| 28.57 | GD | GD | 10.08±8.75 \| 42.86 |
| *Isotricha intestinalis* | 8.96±10.68 \| 100 | 34.50±30.09 \| 100 | GD | 22.42±30.75 \| 100 |
| *Isotricha paraprostoma* | 0.96±0.69 \| 85.71 | 2.06±1.66 \| 100 | GD | 0.81±0.56 \| 85.71 |
| *Isotricha prostoma* | 1.03±0.69 \| 85.71 | 2.75±2.59 \| 100 | 1.01±0.77 \| 100 | 1.01±0.59 \| 85.71 |
| *Ophryoscolex caudatus* | ND | ND | 1.08±0.82 \| 25 | ND |
| *Ostracodinium gracile* | 4.44±5.01 \| 100 | 0.98±0.97 \| 87.50 | 1.98±1.98 \| 87.50 | 1.34±1.17 \| 100 |
| *Polyplastron multivesiculatum* | 21.94±30.11 \| 85.71 | 12.08±18.73 \| 87.50 | 26.36±28.70 \| 87.50 | 17.53±23.26 \| 85.71 |

**Note:** NA, no sequence available in the SILVA or NCBI ITS RefSeq databases representing the respective species; ND, not detected by the respective marker; GD, the species was not detected, but the corresponding genus was detected.

In the rumen samples, only the class Litostomatea was detected and identified (Fig. 5), which is not surprising as all the rumen ciliate detected heretofore have been classified to this class. The four markers consistently detected similar numbers of orders, families, genera, and species of rumen ciliates (Table 4). However, ITS1 and the 28S rRNA gene yielded more ASVs and one additional species compared to the 18S rRNA gene. The RCROD database contains entries representing 17 ciliate species within 12 genera sampled from dairy and beef cattle and goats. Collectively, the four markers detected 13 species within nine genera from the rumen samples collected from dairy cows. Only several species were detected by all the markers, and all the species detected by the 18S rRNA gene were detected by the alternative markers. These results thus show that the alternative markers can perform better than the 18S rRNA gene in detecting and identifying ciliates in freshwater and the rumen ecosystem.

**Fig. 5.**
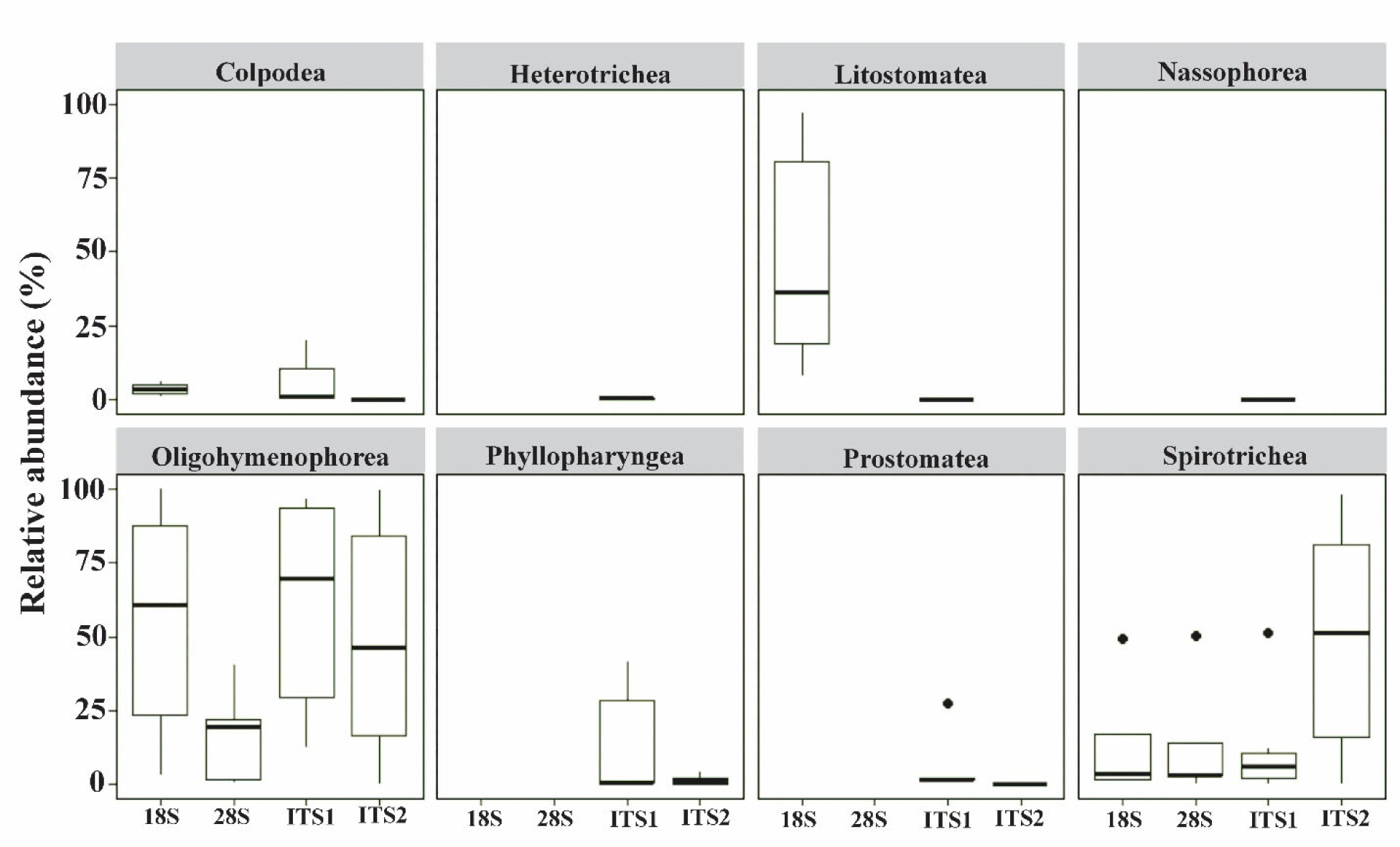
Relative abundance of classes detected by the four phylogenetic markers in freshwater samples.

#### Predominance and prevalence of ciliate classes and species detected in the freshwater and rumen samples

We computed the relative abundance as a measure of predominance at the class level, a taxonomic rank at which morphological features serve as useful criteria for classification. Within the freshwater samples, eight classes of ciliates were detected and identified. The predominance of freshwater ciliates varied considerably depending on the marker used. Heterotrichea and Nassophorea were detected only by the ITS1, while Phyllopharyngea and Prostomatea were only detected by ITS1 and ITS2. Among the classes detected by two or more alternative markers and the 18S rRNA gene, Colpodea and Spirotrichea were detected at comparable predominance, except for Spirotrichea being detected at higher predominance by ITS2. Oligohymenophorea was detected by all the markers as the most predominant class, with its predominance reaching approximately 70%. ITS1 detected it at a higher predominance, but ITS2 and the 28S rRNA gene detected it at a lower predominance. Species belonging to Oligohymenophorea and Spirotrichea, including *Paramecium caudatum, P. multimicronucleatum, Protocyclidium citrillus, Tetrahymena glochidiophila, Strombidinopsis batos, Stylonychia mytilus,* and *S. pustulata*, were detected at high predominance and/or prevalence by ITS1 and ITS2 (Table 5). This aligns with the high prevalence of these two ciliate classes in freshwater ecosystems [60–64]. These results indicate that both ITS markers can support metataxonomic analysis of these two ciliate classes in freshwater ecosystems. At the species level, the predominance of the freshwater ciliates detected and identified by the four markers also varied considerably (Table 5). The 18S rRNA gene detected *Enchelys gasterosteus* of Litostomatea as the most predominant (>80%) species, while *Paramecium micromacronucleatum* of Oligohymenophorea and *Stylonychia pustulata* of Spirotrichea were detected as the most predominant species by ITS1, and ITS2, respectively. The discrepancy in the predominance of ciliates among the markers can be attributed to many factors, including the number of species detected.

Many ciliate species were consistently detected in the rumen samples by the alternative markers (Table 5). *Entodinium caudatum*, the most prevalent species of rumen ciliates, was detected by all four phylogenetic markers across all the samples. Additionally, some species, such as *Entodinium bursa, E. caudatum, E. longinuclatum, Isotricha intestinalis, I. paraprostoma,* and *I. prostoma* were detected across most of the rumen samples. However, different rumen ciliate species were detected as the most predominant species by the four markers (Table 5). *Isotricha intestinalis* was detected as the most predominant species by both ITS1 and the 28S rRNA gene, while *Epidinium caudatum* and *Polyplastron multivesiculatum* were detected as the most predominant species by the 18S rRNA gene and ITS2, respectively. However, *Epidinium caudatum* was not detected by ITS1 or ITS2 due to its absence in the RCTOD database. More SAGs (more than 100) of rumen ciliates are being sequenced (unpublished data), and the ROCRD database will be expanded to include the marker sequences of these new SAGs to further enhance the metataxonomic analysis of the rumen ciliate communities.

Pielou’s evenness was calculated for the four markers to examine the equitability of species abundance distribution within the ciliate communities in the freshwater and rumen samples. For the freshwater ciliate communities, ITS1 and the 28S rRNA gene resulted in a numerically higher evenness than the 18S rRNA gene and ITS2 (Table 4). In the rumen ciliates, the 28S rRNA gene and ITS2 yielded significantly higher evenness values, while ITS1 significantly exhibited a lower evenness value than the 18S rRNA gene. The variations in evenness reflect the variations in the number of species detected and their predominance.

The detection and identification of microbial species in a microbiome through metataxonomics depends on several factors. These include the taxonomic resolution of the phylogenetic marker, the inclusiveness of the primers targeting the marker, potential PCR bias, the comprehensiveness of the database, and the taxonomy implemented in the database. As discussed above, the lack of detection of specific taxa by a marker could be due to the unavailability of the corresponding sequences of those markers in the database used for taxonomic classification. This is particularly true for the 18S and the 28S rRNA genes of ciliates in the current SILVA database. Additionally, the comprehensiveness of the database can also affect the predominance and evenness of the detected ciliate taxa. An artificial mock community of freshwater ciliates and rumen ciliates, with known species composition and predominance, would have allowed better evaluation of the alternative marker in comparison with the 18S rRNA gene, but no such mock ciliate community is available. Thus, we evaluated the four markers and their primers using the same sets of freshwater and rumen samples, focusing on comparison between the two ITS markers and the two rRNA genes. As observed in fugal metataxonomic analysis [65–67], our evaluations and comparisons demonstrate that ITS1, followed by ITS2, resolved more ASVs and detected and identified more species, and higher taxa, in both freshwater and rumen ciliate communities (Tables 4 and 5). ITS1, to a lesser extent ITS2, can be used to enhance metataxonomic analysis of free-living ciliates and rumen ciliates. It should be noted that the macronuclei of ciliates each contain a large copy number of rRNA gene operons. For instance, the genome of *Entodinium caudatum* has 1,010 to 6,210 gene copies [68], and the genome of *Tetrahymena thermophila* has approximately 9,000 gene copies [69]. Therefore, both ITS markers can be well represented in metagenomic sequences and they can be used to facilitate detection and identification of ciliates from metagenomic sequences.

## Conclusions

Metataxonomics remains an essential technology for comprehensively analyzing ciliate communities. Our study assessed the sequence similarity and taxonomic resolution of four phylogenetic markers, including ITS1, ITS2, the 18S, and the 28S rRNA genes, designed and verified primers specific to each marker, and subsequently evaluated their capability in metataxonomic analysis of free-living ciliates in freshwater samples and host-associated ciliates within the rumen ecosystem. Our investigation revealed that ITS1, and to a lesser extent ITS2, offers superior taxonomic resolution compared to the two rRNA genes and enhances comprehensive analysis of ciliate communities in freshwater and rumen ecosystems through metataxonomics. Notably, ITS1 enabled the detection and identification of a broader range of taxa, from class to species, including distinct ciliate species, in both ecosystems. Besides, comparisons of the SILVA and the NCBI ITS RefSeq databases showed that the latter contains more classified taxa, making it more useful in metataxonomic analyses of ciliate species. However, ITS1 failed to detect a small number of the free-living ciliate species detected by the 18S rRNA gene in the freshwater samples due to the absence of the corresponding ITS1 sequences in the NCBI ITS RefSeq database. As the NCBI ITS RefSeq database continues to expand, the capability and utility of ITS1 in comprehensive metataxonomic analyses of ciliate communities in various microbiomes will improve.

## Declarations

### Ethics approval and consent to participate

Not applicable

### Consent for publication

Not applicable

### Availability of data and material

The rumen ciliate rRNA operon database (RCROD) for the 18S, ITS1-5.8S-ITS2, and the 28S rRNA genes are deposited in NCBI GenBank (GenBank submission number: PP067594 to PP067647).

## Authors’ contributions

SS and ZT conceived and designed the study. SS performed the experiment and bioinformatic analyses. SS and ZT wrote and edited the manuscript.

## Acknowledgments

The authors gratefully acknowledge the Molecular and Cellular Imaging Center (MCIC), The Ohio State University, for performing metagenomic sequencing using the Illumina MiSeq system.

## References

1. Gao F, Warren A, Zhang Q, Gong J, Miao M, Sun P, et al. The All-Data-Based evolutionary hypothesis of ciliated protists with a revised classification of the Phylum Ciliophora (Eukaryota, Alveolata). Sci Rep. 2016;6:24874.

2. Dziallas C, Allgaier M, Monaghan MT, Grossart H-P. Act together—implications of symbioses in aquatic ciliates. Front Microbiol. 2012;3.

3. Weisse T. Functional diversity of aquatic ciliates. Eur J Protistol. 2017;61:331–58.

4. Dopheide A, Lear G, Stott R, Lewis G. Relative diversity and community structure of ciliates in stream biofilms according to molecular and microscopy methods. Appl Environ Microbiol. 2009;75:5261–72.

5. Adl SM, Bass D, Lane CE, Lukeš J, Schoch CL, Smirnov A, et al. Revisions to the classification, nomenclature, and diversity of eukaryotes. J Eukaryot Microbiol. 2019;66:4–119.

6. Newbold CJ, De La Fuente G, Belanche A, Ramos-Morales E, McEwan NR. The role of ciliate protozoa in the rumen. Front Microbiol. 2015;6.

7. Bjorbækmo MFM, Evenstad A, Røsæg LL, Krabberød AK, Logares R. The planktonic protist interactome: where do we stand after a century of research? ISME J. 2020;14:544– 59.

8. Zhao Y, Langlois GA. Ciliate morpho-taxonomy and practical considerations before deploying metabarcoding to ciliate community diversity surveys in urban receiving waters. Microorganisms. 2022;10:2512.

9. Warren A, Patterson DJ, Dunthorn M, Clamp JC, Achilles-Day UEM, Aescht E, et al. Beyond the “Code”: a guide to the description and documentation of biodiversity in ciliated protists (Alveolata, Ciliophora). J Eukaryot Microbiol. 2017;64:539–54.

10. Song Y, Hao T, Li B, Zheng W, Liu L, Wang L, et al. Study on analysis of several molecular identification methods for ciliates of Colpodea (Protista, Ciliophora). Cell. Microbiol. 2022;2022:4017442.

11. Keller M, Zengler K. Tapping into microbial diversity. Nat Rev Microbiol. 2004;2:141– 50.

12. Karnati SKR, Yu Z, Sylvester JT, Dehority BA, Morrison M, Firkins JL. Technical note: specific PCR amplification of protozoan 18S rDNA sequences from DNA extracted from ruminal samples of cows. J Anim Sci. 2003;81:812–5.

13. Clemmons BA, Shin SB, Smith TPL, Embree MM, Voy BH, Schneider LG, et al. Ruminal protozoal populations of Angus Steers differing in feed efficiency. Animals (Basel). 2021;11:1561.

14. Shazib SUA, Vd’açny P, Slovák M, Gentekaki E, Shin MK. Deciphering phylogenetic relationships and delimiting species boundaries using a Bayesian coalescent approach in protists: a case study of the ciliate genus *Spirostomum* (Ciliophora, Heterotrichea). Sci Rep. 2019;9:16360.

15. Stern R, Kraberg A, Bresnan E, Kooistra WHCF, Lovejoy C, Montresor M, et al. Molecular analyses of protists in long-term observation programmes—current status and future perspectives. J Plankton Res. 2018;40:519–36.

16. Forster D, Filker S, Kochems R, Breiner H-W, Cordier T, Pawlowski J, et al. A comparison of different ciliate metabarcode genes as bioindicators for environmental impact assessments of Salmon aquaculture. J Eukaryot Microbiol. 2019;66:294–308.

17. Gao F, Kartz LA, Song W. Insights into the phylogenetic and taxonomy of philasterid ciliates (Protozoa, Ciliophora, Scuticociliata) based on analyses of multiple molecular markers. Mol Phylogenet Evol. 2012;64:308–17.

18. Wang C, Hu Y, Warren A, Hu X. Genetic diversity and phylogeny of the genus *Euplotes* (Protozoa, Ciliophora) revealed by the mitochondrial CO1 and nuclear ribosomal genes. Microorganisms 2021;9:2204.

19. Zhao Y, Yi Z, Gentakaki E, Zhan A, Al-Farraj SA, Song W. Utility of combining morphological characters, nuclear and mitochondrial genes: an attempt to resolve the conflicts of species identification for ciliated protists. Mol Phylogenet Evol. 2016;94:718– 29.

20. Zhang Q, Xu J, Warren A, Yang R, Shen Z, Yi Z. Assessing the utility of Hsp90 gene for inferring evolutionary relationships within the ciliate subclass Hypotrichia (Protista, Ciliophora). BMC Evol Biol. 2020;20:86.

21. Bachy C, Dolan JR, López-García P, Deschamps P, Moreira D. Accuracy of protist diversity assessments: morphology compared with cloning and direct pyrosequencing of 18S rRNA genes and ITS regions using the conspicuous tintinnid ciliates as a case study. The ISME J. 2013;7:244–55.

22. Li J, Liu W, Gao S, Warren A, Song W. Multigene-based analyses of the phylogenetic evolution of oligotrich ciliates, with consideration of the internal transcribed spacer 2 secondary structure of three systematically ambiguous genera. Eukaryot Cell. 2013;12:430–7.

23. Miao M, Warren A, Song W, Wang S, Shang H, Chen Z. Analysis of the internal transcribed spacer 2 (ITS2) region of Scuticociliates and related taxa (Ciliophora, Oligohymenophorea) to infer their evolution and phylogeny. Protist 2008;159:519–33.

24. Obert T, Vďačný, P. Delimitation of five astome ciliate species isolated from the digestive tube of three ecologically different groups of lumbricid earthworms, using the internal transcribed spacer region and the hypervariable D1/D2 region of the 28S rRNA gene. BMC Evol Biol. 2020;20:37.

25. Ogier J-C, Pagès S, Galan M, Barret M, Gaudriault S. rpoB, a promising marker for analyzing the diversity of bacterial communities by amplicon sequencing. BMC Microbiol. 2019;19:171.

26. Li L, Meng D, Yin H, Zhang T, Liu Y. Genome-resolved metagenomics provides insights into the ecological roles of the keystone taxa in heavy-metal-contaminated soils. Front Microbiol. 2023;14:1203164.

27. Reji L, Cardarelli EL, Boye K, Bargar JR, Francis CA. Diverse ecophysiological adaptations of subsurface Thaumarchaeota in floodplain sediments revealed through genome-resolved metagenomics. ISME J. 2022;16:1140–52.

28. Stewart RD, Auffret MD, Warr A, Walker AW, Roehe R, Watson M. Compendium of 4,941 rumen metagenome-assembled genomes for rumen microbiome biology and enzyme discovery. Nat Biotechnol. 2019;37:953–61.

29. Blaalid R, Kumar S, Nilsson RH, Abarenkov K, Kirk PM, Kauserud H. ITS1 versus ITS2 as DNA metabarcodes for fungi. Mol Ecol Resour. 2013;13:218–24.

30. Heeger F, Wurzbacher C, Bourne EC, Mazzoni CJ, Monaghan MT. Combining the 5.8S and ITS2 to improve classification of fungi. Methods Ecol Evol. 2019;10:1702–11.

31. Li B, Song Y, Hao T, Wang L, Zheng W, Lyu Z, et al. Insights into the phylogeny of the ciliate of class Colpodea based on multigene data. Ecol Evol. 2022;12:e9380.

32. Liu L, Jiang M, Zhou C, Li B, Song Y, Pan X. Further insights into the phylogeny of facultative parasitic ciliates associated with tetrahymenosis (Ciliophora, Oligohymenophorea) based on multigene data. Ecol Evol. 2023;13:e10504.

33. Liao W, Gong Z, Ni B, Fan X, Petroni G. Documentation of a new hypotrich species in the family Amphisiellidae, *Lamtostyla gui* n. sp. (Protista, Ciliophora) using a multidisciplinary approach. Sci Rep. 2020;10:3763.

34. Frantal D, Agatha S, Beisser D, Boenigk J, Darienko T, Dirren-Pitsch G, et al. Molecular data reveal a cryptic diversity in the genus *Urotricha* (Alveolata, Ciliophora, Prostomatida), a key player in freshwater lakes, with remarks on morphology, food preferences, and distribution. Front Microbiol. 2021;12:787290.

35. Zhou T, Wang Z, Yang H, Gu Z. Two new colonial peritrich ciliates (Ciliophora, Peritrichia, Sessilida) from China: With a note on taxonomic distinction between Epistylididae and Operculariidae. Eur J Protistol. 2019;70:17–31.

36. Robeson MS, O’Rourke DR, Kaehler BD, Ziemski M, Dillon MR, Foster JT, et al. RESCRIPt: Reproducible sequence taxonomy reference database management. PLoS Comput Biol. 2021;17:e1009581.

37. Li Z, Wang X, Zhang Y, Yu Z, Zhang T, Dai X, et al. Genomic insights into the phylogeny and biomass-degrading enzymes of rumen ciliates. ISME J. 2022;16:2775–87.

38. Tamura K, Stecher G, Kumar S. MEGA11: Molecular Evolutionary Genetics Analysis Version 11. Molecular Biology and Evolution. 2021;38:3022–7.

39. Hall TA. BioEdit: a user-friendly biological sequence alignment editor and analysis program for Windows 95/98/NT. Nucleic Acids Symp Ser. 1999;41:95–8.

40. Untergasser A, Cutcutache I, Koressaar T, Ye J, Faircloth BC, Remm M, et al. Primer3— new capabilities and interfaces. Nucleic Acids Res. 2012;40:e115.

41. Sylvester JT, Karnati SKR, Yu Z, Morrison M, Firkins JL. Development of an assay to quantify rumen ciliate protozoal biomass in cows using real-time PCR. J Nutr. 2004;134:3378–3384.

42. Ishaq SL, Wright A-DG. Design and validation of four new primers for next-generation sequencing to target the 18S rRNA genes of gastrointestinal ciliate protozoa. Appl Environ Microbiol. 2014;80:5515–21.

43. Kalendar R. A guide to using FASTPCR software for PCR, in silico PCR, and oligonucleotide analysis. In: Basu C, editor. PCR Primer Design. New York. Springer; 2022. p. 223–43.

44. Larkin MA, Blackshields G, Brown NP, Chenna R, McGettigan PA, McWilliam H, Valentin F, Wallace IM, Wilm A, Lopez R, Thompson JD, Gibson TJ, Higgins DG. Clustal W and Clustal X version 2.0. Bioinformatics. 2007;23;2947–8.

45. Tamura K. Estimation of the number of nucleotide substitutions when there are strong transition-transversion and G + C-content biases. Mol Biol Evol. 1992;9:678–87.

46. Bolyen E, Rideout JR, Dillon MR, Bokulich NA, Abnet CC, Al-Ghalith GA, et al. Reproducible, interactive, scalable and extensible microbiome data science using QIIME 2. Nat Biotechnol. 2019;37:852–7.

47. McMurdie PJ, Holmes S. phyloseq: An R Package for Reproducible Interactive Analysis and Graphics of Microbiome Census Data. PLoS ONE. 2013;8:e61217.

48. Gupta R, Abraham JS, Sripoorna S, Maurya S, Toteja R, Makhija S, et al. Description of a new species of *Tetmemena* (Ciliophora, Oxytrichidae) using classical and molecular markers. J King Saud Univ Sci. 2020.32:2316–28.

49. Park T, Meulia T, Firkins JL, Yu Z. Inhibition of the rumen ciliate *Entodinium caudatum* by antibiotics. Front Microbiol. 2017;8:1189.

50. R Core Team. R: A language and environment for statistical computing. Vienna, Austria: R foundation for statistical computing. 2023. https://www.R-project.org/

51. Wickham H. ggplot2: Elegant Graphics for Data Analysis. New York: Springer-Verlag; 2016.

52. Wickham H, Averick M, Bryan J, Chang W, McGowan LDA, François R, et al. (2019). Welcome to the Tidyverse. J Open Source Softw. 2019;4:1686.

53. Oksanen J, Simpson GL, Blanchet FG, Kindt R, Legendre P, Minchin PR, et al. vegan: community ecology package. 2022. https://github.com/vegandevs/vegan

54. Zhan Z, Li J, Xu K. Ciliate environmental diversity can be underestimated by the V4 region of SSU rDNA: insights from species delimitation and multilocus phylogeny of *Pseudokeronopsis* (Protist, Ciliophora). Microorganisms 2019;7:493.

55. Wright ADG, Lynn DH. Monophyly of the Trichostome ciliates (Phylum Ciliophora: Class Litostomatea) tested using new 18S rRNA sequences from the Vestibuliferids, *Isotricha intestinalis* and *Dasytricha ruminantium*, and the Haptorian, *Didinium nasutum*. Eur J Protistol 1997;33:305–15.

56. Da Silva ZRJ, Cedrola F, Rossi MF, Costa FDS, Dias RJP. Rumen ciliates (Alveolata, Ciliophora) associated with goats: checklist, geographic distribution, host specificity, phylogeny and molecular dating. Zootaxa 2022;5165:2.

57. Chan AHE, Saralamba N, Saralamba S, Ruangsittichai J, Urusa T. The potential use of mitochondrial ribosomal genes (12S and 16S) in DNA barcoding and phylogenetic analysis of trematodes. BMC Genomics 2022;23:104.

58. Kounosu A, Murase K, Yoshida A, Maruyama H, Kikuchi T. Improved 18S and 28S rDNA primer sets for NGS-based parasite detection. Sci Rep. 2019;9:15789.

59. Hillis DM, Dixon MT. Ribosomal DNA: molecular evolution and phylogenetic inference. Q Rev Biol. 1991;66,:411–53.

60. Abraham JS, Gupta R, Somasundaram S, Naqvi I, Maurya S, Toteja R, et al. Faunistic study on the freshwater ciliates from Delhi, India. https://www.biorxiv.org/content/10.1101/2020.07.06.189001v1.full (2020). Accessed 06 Jul 2020.

61. Dinçer SC. Freshwater ciliates from Beytepe Pond in Ankara with new records for Turkey. Turk J Zool. 2016;40;5.

62. Jiang C, Liu B, Zhang J, Gu S, Liu Z, Wang X, et al. Diversity and seasonality dynamics of ciliate communities in four estuaries of Shenzhen, China (South China Sea). J Mar Sci Eng. 2021;9:260.

63. Sun P, Huang L, Xu D, Huang B, Chen N, Warren A. Marked seasonality and high spatial variation in estuarine ciliates are driven by exchanges between the ‘abundant’ and ‘intermediate’ biospheres. Sci Rep. 2017;7:9494.

64. Sun P, Huang L, Xu D, Warren A, Huang B, Wang Y, et al. Integrated Space-Time Dataset Reveals High Diversity and Distinct Community Structure of Ciliates in Mesopelagic Waters of the Northern South China Sea. Front Microbiol. 2019;10:2178.

65. Hoggard M, Vesty A, Wong G, Montgomery JM, Fourie C, Douglas RG, et al. Characterizing the human mycobiota: a comparison of small subunit rRNA, ITS1, ITS2, and large subunit rRNA genomic targets. Front Microbiol. 2018;9:2208.

66. Mbareche H, Veillette M, Bilodeau G, Duchaine C. Comparison of the performance of ITS1 and ITS2 as barcodes in amplicon-based sequencing of bioaerosols. PeerJ. 2020;8:e8523.

67. Wang X-C, Liu C, Huang L, Bengtsson-Palme J, Chen H, Zhang J-H, et al. ITS1: a DNA barcode better than ITS2 in eukaryotes? Mol Ecol Resour. 2015;15:573–86.

68. Sylvester JT, Karnati SK, Yu Z, Newbold CJ, Firkins JL. Evaluation of a real-time PCR assay quantifying the ruminal pool size and duodenal flow of protozoal nitrogen. J Dairy Sci. 2005;88:2083–95.

69. Blomberg P, Randolph C, Yao CH, Yao MC. Regulatory sequences for the amplification and replication of the ribosomal DNA minichromosome in *Tetrahymena thermophila*. Mol Cell Biol. 1997;17:7237–47.

