## Supplemental Table 1 for "Internal Transcribed Spacers as Phylogenetic Markers Enable Species-level Metataxonomic Analysis of Ciliated Protozoa"

**Table S1** Contigs of rumen ciliates containing ribosomal rRNA genes and ITS regions.

| Species name | Contigs | 18S rRNA gene | ITS region | 28S rRNA gene |
|---|---|---|---|---|
| | | **Location (Length) in bp** | | |
| *Dasytricha ruminantium* | SAG2_0_NT_8340 | 1,004-1,483 (480) | 2,308-2,761 (454) | 3,049-3,325 (277) |
| | SAG3_0_ST_7869 | 2,574-3,053 (480) | 3,878-4,331 (454) | 4,619-4,895 (277) |
| | SAG4_0_ST_7428 | 2,180-2,659 (480) | 3,484-3,937 (454) | 4,225-4,501 (277) |
| | SAG5_0_NT_10536 | 5,302-5,781 (480) | 6,606-7,060 (455) | 7,348-7,624 (277) |
| | SAGT1_0_NT_5350 | 279-758 (480) | 1,583-2,036 (454) | 2,324-2,600 (277) |
| *Diplodinium dentatum* | SAG3_0_NT_5349 | 316-796 (481) | 1,62-2,094 (474) | 2,382-2,658 (277) |
| | SAG4_0_ST_8575 | 3,540-4,020 (481) | 4,845-5,318 (474) | 5,606-5,882 (277) |
| | SAGT1_0_NT_5669 | 636-1,116 bp (481) | 1,941-2,414 (474) | 2,702-2,978 (277) |
| | SAGT2_0_NT_8265 | 3,145-3,625 (481) | 4,450-4,923 (474) | 5,211-5,487 (277) |
| *Diplodinium flabellum* | SAG1_0_BT_9736 | 4,344-4,824 (481) | 5,649-6,121 (473) | 6,409-6,685 (277) |
| | SAG2_0_NT_7877 | 2,107-2,587 (481) | 3,412-3,884 (473) | 4,172-4,448 (277) |
| *Enoploplastron triloricatum* | SAG3_0_ST_9549 | 3,194-3,674 (481) | 4,499-4,971 (473) | 5,259-5,535 (277) |
| | SAGT1_0_NT_6595 | 1,348-1,828 (481) | 2,653-3,125 (473) | 3,413-3,689 (277) |
| | SAGT2_0_ST_9012 | 3,774-4,254 (481) | 5,097-5,551 (455) | 5,839-6,115 (277) |
| *Entodinium bursa* | SAG3_0_NT_6796 | 1,499-1,980 (482) | 2805-3,282 (478) | 3,570-3,844 (275) |
| | SAGT1_0_BT_9368 | 4,133-4,647 (482) | 5,439-5,916 (478) | 6,204-6,478 (275) |
| *Entodinium caudatum* | SAG3_0_NT_6493 | 1,365-1,846 (482) | 2,671-3,145 (475) | 3,433-3,707 (275) |
| | SAG4_0_BT_10558 | 4,031-4,512 (482) | 5,337-5,811 (475) | 6,099-6,373 (275) |
| | SAGT1_0_ST_7247 | 2,055-2,536 (482) | 3,361-3,835 (475) | 4,123-4,397 (275) |
| *Entodinium longinucleatum* | SAGT1_0_BT_8110 | 2,937-3,418 (482) | 4,243-4,721 (479) | 5,009-5,275 (267) |
| | SAG4_0_BT_8110 | 2,926-3,407 (482) | 4,232-4,710 (479) | 4,998-5,264 (267) |
| | SAGT2_0_ST_8067 | 2,919-3,400 (482) | 4,225-4,703 (479) | 4,991-5,257 (267) |

| | | | | |
|---|---|---|---|---|
| | SAGT3_0_NT_5310 | 201-682 (482) | 1,507-1,985 (479) | 2,273-2,539 (267) |
| *Epidinium cattanei* | SAG2_0_NT_7670 | 2,244-2,723 (480) | 3,550-4,023 (474) | 4,311-4,586 (276) |
| | SAG3_0_ST_8014 | 2728-3,207 (480) | 4,034-4,507 (474) | 4,795-5,070 (276) |
| *Epidinium caudatum* | SAG1_0_BT_13234 | 2,769-3,248 (480) | 4,075-4,548 (474) | 4,836-5,112 (277) |
| | SAG3_0_NT_7623 | 2,368-2,847 (480) | 3,674-4,147 (474) | 4,435-4,711 (277) |
| *Eremoplastron rostratum* | SAG1_0_ST_8555 | 3,271-3,750 (480) | 4,574-5,048 (475) | 5,336-5,612 (277) |
| | SAG2_0_NT_8259 | 3,043-3,522 (480) | 4,346-4,820 (475) | 5,120-5,384 (265) |
| *Isotricha intestinalis* | SAG3_0_NT_56080 | 14,188-14,670 (483) | 15,497-15,947 (451) | 16,236-16,512 (277) |
| | SAGT1_0_NT_14931 | 7,901 to 8,383 (483) | 9,210-9,660 (451) | 9,960-10,225 (266) |
| | SAGT2_0_ST_9665 | 2,746-3,228 (483) | 4,055-4,505 (451) | 4,794-5,070 (277) |
| *Isotricha jalaludinii* | SAG1_0_ST_15841 | 2,458-2,941 (484) | 3,767-4,216 (450) | 4,504-4,780 (277) |
| | SAG2_0_NT_16005 | 2,598-3,081 (484) | 3,907-4,356 (450) | 4,644-4,920 (277) |
| | SAG4_0_NT_119093 | 105,794-106,277 (484) | 107,103-107,552 (450) | 107,840-108,116 (277) |
| | SAG5_0_NT_16511 | 3,031-3,514 (484) | 4,340-4,789 (450) | 5,077-5,353 (277) |
| *Isotricha paraprostoma* | SAG1_0_BT_7142 | 1,894-2,376 (483) | 3,201-3,647 (447) | 3,934-4,210 (277) |
| *Isotricha prostoma* | SAG2_0_ST_7329 | 1,940-2,423 (484) | 3,248-3,693 (446) | 3,981-4,257 (277) |
| | SAGT1_0_NT_9389 | 3,642-4,125 (484) | 4.950-5,395 (446) | 5,683-5,959 (277) |
| *Metadinium minomm* | SAG1_0_ST_7813 | 2,789-3,270 (482) | 4,095 to 4,567 (473) | 4,855-5,131 (277) |
| *Ophryoscolex caudatus* | SAG5_0_BT_8594 | 2,601-3,080 (480) | 3,905 to 4,379 (475) | 4,667-4,942 (276) |
| | SAG6_0_ST_7845 | 2,568-3,047 (480) | 3,872-4,346 (475) | 4,634-4,909 (276) |
| | SAG8_0_ST_7556 | 2,287-2,766 (480) | 3,591-4,069 (479) | 4,353-4,628 (276) |
| | SAG11_0_NT_6766 | 1,733-2,212 (480) | 3,037-3,511 (475) | 3,799-4,074 (276) |
| | SAGT3_0_BT_9281 | 4,023-4,502 (480) | 5,327-5,801 (475) | 6,089-6,364 (275) |
| | SAGT4_0_NT_8124 | 2,895-3,374 (480) | 4,199-4,673 (475) | 4,961-5,236 (276) |
| *Ostracodinium dentatum* | SAG1_0_BT_8444 | 2,711-3,190 (480) | 4,016-4,487 (472) | 4,775-5,051 (277) |

| | | | | |
|---|---|---|---|---|
| *Ostracodinium gracile* | SAG1_0_NT_6491 | 1,483-1,963 (481) | 2,789-3,257 (469) | 3,545-3,821 (277) |
| | SAG2_0_NT_5520 | 328-808 (481) | 1,634-2,102 (468) | 2,390-2,666 (277) |
| | SAG3_0_ST_8191 | 3,004-3,484 (481) | 4,310-4,778 (469) | 5,066-5,342 (277) |
| *Polyplastron multivesiculatum* | SAG8_0_ST_7510 | 1,667-2,148 (482) | 2,974-3,447 (474) | 3,736-4,012 (277) |
| | SAGT1_0_NT_6642 | 1,212-1,693 (482) | 2,519-2,992 (474) | 3,281-3,557 (277) |
| | SAGT2_0_NT_7085 | 1245-1,726 (482) | 2,552-3,025 (474) | 3,314-3,590 (277) |
| | SAGT3_0_ST_8853 | 1,487-1,968 (482) | 2,794-3,267 (474) | 3,556-3,832 (277) |
| | SAGT4_0_ST_6441 | 1,212-1,693 (482) | 2,519-2,992 (474) | 3,281-3,557 (277) |
| | SAGT5_0_ST_6296 | 1,081-1,562 (482) | 2,388-2,861 (474) | 3,150-3,426 (277) |
| | SAGT7_0_ST_6430 | 1,212-1,693 (482) | 2,519-2,992 (474) | 3,281-3,557 (277) |
